# Environmental and Maternal Imprints on Infant Gut Metabolic Programming

**DOI:** 10.1101/2025.07.24.666662

**Authors:** Kine Eide Kvitne, Celeste Allaband, Jennifer C. Onuora, Daniela Perry, Simone Zuffa, Lucas Patel, Vincent Charron-Lamoureux, Ipsita Mohanty, Kristija Sejane, Abubaker Patan, Abdullah Al Mahmud, Tahmeed Ahmed, Diego G. Bassani, Antonio González, Davidson H. Hamer, Rashidul Haque, Benjamin Ho, Md Iqbal Hossain, Mohammad Shahidul Islam, Daniel McDonald, Lisa G. Pell, Huma Qamar, Daniel E. Roth, Samir Saha, Prakesh S. Shah, Md Muniruzzaman Siddiqui, Shafiqul Alam Sarker, Shamima Sultana, Sydney Thomas, Lindsey A. Burnett, Shirley M. Tsunoda, Lars Bode, Pieter C. Dorrestein, Rob Knight

**Author notes:** Contributed equally (Kine Eide Kvitne, Celeste Allaband, Jennifer C. Onuora). Correspondence (R.K.; microbiome analysis), and (P.C.D.; metabolomics analysis).

## Abstract

Early life is a critical period for immune and metabolic programming, but developmental patterns remain underexplored in populations from low- and middle-income countries. Here, we profiled the microbiome and metabolome of 55 Bangladeshi mother-infant dyads over the first six months of life. Importantly, we observed an increase in microbially-derived bile amidates and *N*-acyl lipids with age in conjunction with reads matching the bile salt hydrolase/transferase (*bsh*) gene. While microbial source tracking confirmed maternal fecal seeding, a substantial environmental contribution was also highlighted. Differences in infant fecal metabolic profiles were associated with delivery mode, maternal milk composition, household assets, and household-level water treatment. C-section delivery and untreated drinking water were linked to transient metabolic differences, including increases in bile amidates, *N*-acyl lipids, and other host-microbe co-metabolic products, including acylcarnitines. Multi-omics analysis revealed specific microbial-metabolite relationships, highlighting how early environmental and maternal living circumstances shape metabolic gut programming through the microbiome.

## MAIN

The first months after birth represent a crucial window for infant development, laying the groundwork for long-term health. During this period, the establishment and maturation of the gut microbiome play a key role in metabolic programming and training of the infant immune system^1,2^. Although these processes are known to be shaped by a complex interplay between maternal and environmental factors^3,4^, our current knowledge largely originates from studies in high-income countries^5^. Thus, early-life gut microbiome composition and function in low- and middle-income countries (LMICs) largely remain underexplored.

Delivery mode is one of the earliest determinants of newborn microbiota colonization^6,7^. While vaginally born infants are seeded by maternal fecal and vaginal microbiota, infants born via Cesarean section (C-section) are primarily colonized by skin associated microorganisms, such as *Staphylococcus* and *Corynebacterium^6,8^*. C-section births have increased by nearly 30% over the last decade in many LMICs, including Bangladesh^9^. After birth, challenges with early initiation and duration of breastfeeding^10–12^ can limit infant exposure to bioactive human milk oligosaccharides (HMOs). These molecules function as prebiotics, fostering the growth of the infant gut microbiota and influencing health outcomes^13,14^. For example, α1-2-fucosylated HMOs, like 2’-fucosyllactose (2’FL), produced almost exclusively by mothers with a functional fucosyltransferase 2 (FUT2) gene^15^ (Secretors) can promote early-life growth of beneficial microorganisms, such as *Bifidobacterium^16,17^*. In contrast, the abundance of *Bifidobacterium* species is negatively impacted by antibiotic treatment in early infancy^18^, along with overall changes to microbiome diversity and composition^7,19,20^. Antibiotic usage is considerably more common in many LMICs^21–23^. In addition, variable household hygiene standards and sanitation practices^24^ which are generally connected to wealth may further influence the microbiome development through different exposure to pathobionts or molecular exposures.

Since infant microbiomes do not function in isolation, integrating microbiome and metabolomics data through multi-omics approaches is essential to understand how early-life exposures and living conditions shape gut metabolic programming. However, such efforts have been limited by the technical difficulty of acquiring rich, longitudinal multi-omics data, particularly in LMIC settings. Here, we address this gap through an observational, longitudinal study of 55 mother-infant dyads in Bangladesh, integrating metagenomics, untargeted metabolomics, and HMO profiling across the first six months of life. Our study demonstrates a rapid increase in microbially-derived metabolites, including bile amidates and *N*-acyl lipids, in response to the maturing gut microbiome. We also show that delivery mode, maternal secretor status, and household drinking water treatment are associated with different infant fecal metabolomic profiles. Finally, multi-omics integration reveals specific microbial-metabolite relationships, highlighting the role of the gut microbiome as a key mediator of early metabolic programming.

## RESULTS

We enrolled 55 mother-infant dyads from two urban hospitals in Dhaka, Bangladesh, and sampled them at multiple timepoints from birth to six months postnatal age **(Fig. 1a, detailed flow chart in Extended Data Fig. 1a)**. Infants between 0-1 days old and orally feeding at the time of screening were eligible for inclusion. We collected stool samples from both infants and mothers, human milk and vaginal swabs from mothers, and blood, skin (elbow), and oral cavity (tongue) samples from infants. Untargeted liquid chromatography coupled with tandem mass spectrometry (LC-MS/MS) metabolomics was used to analyze infant stools, plasma, and maternal milk, while shotgun metagenomic sequencing was performed on all sample types except plasma. A total of 1,609 metagenomics samples and 530 metabolomics samples were processed.

**Fig. 1:**
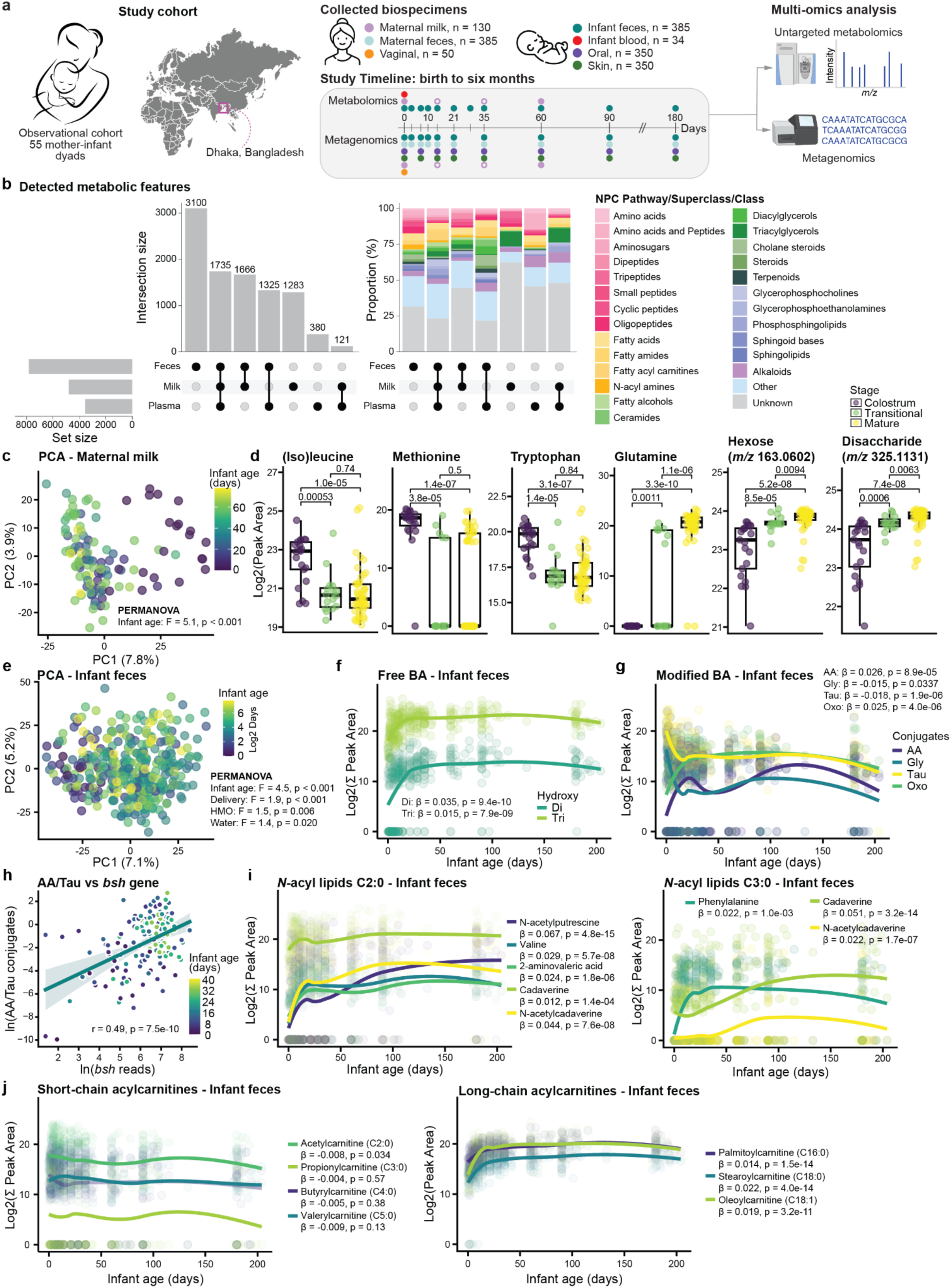
Rapid shifts in key signalling metabolites during infant fecal metabolic gut programming. **a,** Experimental overview. Longitudinal samples were collected from 55 mother-infant dyads in Dhaka, Bangladesh. Infant samples included stool, oral (tongue), skin (elbow), and blood; maternal samples included stool, milk, and a vaginal swab (see Methods for details). Open circles indicate random collection time within the first 60 days. n reflects the initial number of samples per biospecimen, prior to quality control. **b,** UpSet plot of metabolic features across infant fecal (n=377) and plasma (n=34) samples, and maternal milk (n=119) samples. The left panel shows features unique to or shared between sample types, while the right panel shows chemical classes and pathways predictions based on NPC classification from CANOPUS (probability > 0.5). **c,** Principal Component Analysis (PCA) of maternal milk showing significant development over time (Permutational Multivariate Analysis of Variance (PERMANOVA): p < 0.001). **d,** Boxplot showing the natural log peak areas of key discriminant features across lactation stages: colostrum (< 5 days), transitional (5-15 days), and mature milk (> 15 days). Statistical significance was determined by the Wilcoxon rank-sum test, with p values adjusted for multiple comparisons using the Benjamini-Hochberg (BH) method. Repeated measures were avoided by retaining only the highest value per subject within each lactation stage. **e,** PCA of fecal metabolic profiles across early infancy (0-6 months). Differences in the metabolic profiles were associated with infant age, delivery mode, maternal secretor status, and drinking water treatment (PERMANOVA: all p < 0.05). **f,** Longitudinal abundances of free di- and tri-hydroxylated bile acids in infant feces over the first 6 months of life. **g,** Abundance trajectories of taurine (tau), glycine (gly), and amino acid (AA) conjugated bile acids, as well as oxo bile acids in the same period. **h,** Scatter plot showing the association between the natural log of bile salt hydrolase/transferase (*bsh)* gene read count matches and the ratio of amino acid to taurine conjugated bile acids using Pearson correlation for significance (linear mixed effect model (LME): β = 0.25, p < 0.001). A linear regression line with 95% CI is shown, with points colored by infant age. **i,** Longitudinal abundance trajectories of *N*-acyl lipids with a C2:0 fatty acid chain (left panel) and a C3:0 fatty acid chain (right panel) amidated to different head groups. **j,** Longitudinal abundances of MS/MS spectral matches to various short-chain acylcarnitines (left panel) and long-chain acylcarnitines (right panel) in infant feces. For panels **f, g, i, j**, β and p values were derived from LME with subject as a random effect, and scatter plot smoothing (‘LOESS’) was used for visualization purposes. All boxplots show the first (lower), median, and third (upper) quartiles, with whiskers 1.5 times the interquartile range.

Characteristics of the study cohort are detailed in **Table 1**. In short, 40% of infants were male and infants had an average birth weight of 2,924 ± 428 g (mean ± standard deviation). All mothers received postpartum antibiotics, and, in line with the pre-specified enrollment protocol, there was an even distribution between vaginal and C-section deliveries. Exclusive breastfeeding rates declined from 62% at one month to 42% at 3 months to a steep drop of 16% at 6 months. Based on 2’-fucosyllactose (2’FL) levels, 67% of mothers were classified as secretors. Because variable household hygiene standards in LMICs may also influence microbiome composition and function, we also gathered data on variables such as drinking water practices. Importantly, 67% of households reported using at least one method to treat their drinking water (boiling and/or filtering).

**Table 1.** Characteristics of the study cohort. Numbers are given as mean ± SD or count (%).

| Characteristics | Infant (n = 55) |
| --- | --- |
| Infant sex, male | 22 (40%) |
| Birthweight, g | 2,924 ± 428 |
| Gestational age, weeks | 39.1 ± 1.6 |
| Maternal age, years | 26 ± 5 |
| Delivery mode, vaginal | 28 (51%) |
| Feeding patterns |  |
| - Exclusively breastfed ~1 month | 34 (62%) |
| - Exclusively breastfed ~3 months | 23 (42%) |
| - Exclusively breastfed ~6 months | 9 (16%) |
| Secretor status <sup>+</sup> |  |
| - Secretor | 37 (67%) |
| - Non-secretor | 16 (29%) |
| Maternal antibiotics <sup>‡</sup> |  |
| - Intrapartum and postpartum | 13 (24%) |
| - Postpartum only | 42 (76%) |
| Infant antibiotics <sup>#</sup> |  |
| - Ever received by 1 month | 14 (26%) |
| - Ever received by 3 months | 17 (31%) |
| - Ever received by 6 months | 31 (56%) |
| Asset index <sup>*</sup> |  |
| - 1 (lowest wealth) | 6 (11%) |
| - 2 | 9 (16%) |
| - 3 | 6 (11%) |
| - 4 | 18 (33%) |
| - 5 (highest wealth) | 16 (29%) |
| Treatment of drinking water, yes (boiled and/or filtered) <sup>§</sup> | 37 (67%) |

<sup>||</sup>Exclusive breastfeeding is defined as only receiving human milk (no other liquids or solids, including water). Exclusively breastfed status is reported for specific time windows: 1 month (28-35 days), 3 months (89-155 days), and 6 months (177-247 days). The remaining infants in each window were not exclusively breastfed.
<sup>+</sup>Secretors/non-secretors were defined by the presence or near-absence (<100 nmol/mL) of the human milk oligosaccharide 2'-fucosyllactose. Information on secretor status was missing for two subjects.
<sup>‡</sup>The most common antibiotics given to the mothers included metronidazole, ceftriaxone, cephradine, amoxicillin and flucloxacillin.
<sup>#</sup>Reflects the infant's age at first antibiotic exposure. Infants were assigned "Yes" for each time point if the earliest recorded age of antibiotic use fell within the defined window: $\leq 35$ days (1 month), $\leq 90$ days (3 months), or $\leq 180$ days (6 months). The most common antibiotics given to the infants included cefpodoxime proxetil, cefixime, azithromycin, cefaclor, ceftriaxone, amoxicillin, cefradine, gentamicin, and ampicillin.
<sup>\*</sup>Asset index, a proxy for socioeconomic status, was generated as previously described<sup>25</sup> (details in methods) with higher quintiles representing greater wealth and lower quintiles reflecting lower wealth.
<sup>§</sup>Reported to boil and/or filter their drinking water at baseline. The remaining households reported to not apply any form of water treatment.

### Infant Age Shapes Early Metabolic Trajectories in Maternal Milk and Infant Feces

To characterize early-life metabolic trajectories in the mother-infant dyads, we profiled infant feces (n=377), infant plasma (n=34), and maternal milk (n=119) using untargeted LC-MS/MS metabolomics. This resulted in the detection of 13,871 metabolic features, of which 1,735 were shared across all three biospecimens **(Fig. 1b)**. As expected, each sample type displayed a distinct metabolic composition. Maternal milk was rich in amino acids, saccharides, and triacylglycerols, while infant feces had a broader range of metabolite classes, including amino acids, small peptides, fatty acids, *N*-acyl amines, steroids, sphingolipids, and alkaloids.

Maternal milk composition changed over time (permutational multivariate analysis of variance (PERMANOVA): R^2^ = 0.04, F-statistic = 5.1, p < 0.001; **Fig. 1c**), consistent with knowledge that nutritional needs change as the infant grows^26^. To identify metabolic changes across lactation stages, milk samples were grouped into colostrum (< 5 days), transitional (5-15 days), and mature milk (> 15 days), and analysed using a supervised Partial Least Squares Discriminant Analysis (PLS-DA) model (classification error rate (CER) = 0.18; **Extended Data Fig. 2a**). Features with a variable importance in projection (VIP) > 1 were extracted (**Supplementary Table 1**). Colostrum was enriched in amino acids, small peptides, and short-chain acylcarnitines, including acetylcarnitine (C2:0) and propionylcarnitine (C3:0) (**Fig. 1d, Extended Data Fig. 2b**). Here, C3:0 refers to a fatty acid chain with three carbons and no double bonds^27^. In contrast, mature milk contained higher levels of carbohydrate sources, such as mono- and di-saccharides, as well as glutamine, glutamate and butyrylcarnitine (C4:0), consistent with previous findings^26,28^.

We also observed a clear development of the infant fecal metabolome with age (PERMANOVA: R^2^ = 0.012, F-statistic = 4.5, p < 0.001; **Fig. 1e**). Bile acid metabolism, critical for infant development^29^, changed during the first weeks of life. Using recently developed libraries^30–32^, we show that the fecal abundance of free di- and tri-hydroxylated bile acids increased over the first weeks of life (linear mixed effects model (LME): di: β = 0.035, p = 9.4×10^-10^; tri: β = 0.015, p = 7.9×10^-9^; **Fig. 1f**). Host-derived taurine (LME: β = -0.018, p = 1.9×10^-6^) and host-microbially derived glycine (LME: β = -0.015, p = 0.034) conjugates decreased in the same period (**Fig. 1g**). In contrast, microbially modified bile acids, including various amino acid conjugates (LME: β = 0.026, p = 8.9×10^-5^) and oxo bile acids (LME: β = 0.025, p = 4.0×10^-6^) increased (**Fig. 1g**). This opposite trend suggests a growing capacity of the gut microbiome to modify host-derived bile acids as it matures. To investigate this more mechanistically, we looked at the microbial bile salt hydrolase/transferase (*bsh*) gene, which deconjugates taurine and glycine conjugates through hydrolysis of the amide bond but also are capable of conjugating bile acids with amines^33,34^. We observed a positive correlation between *bsh* gene levels and the ratio of amino acid to taurine conjugated bile acids (Pearson correlation: r = 0.49, p = 7.5e-10 ; **Fig. 1h**). Notably, both the abundance of the *bsh* gene family (LME: β = 0.057, p < 0.001; **Extended Data Fig. 2c**) and this bile acid ratio (LME: β = 0.074, p < 0.001; **Extended Data Fig. 2d**) increased with age, linking the maturation of the gut microbiome to the observed shifts in the bile acid pool. Another group of signalling molecules that underwent changes in early life were the *N*-acyl lipids, metabolites consisting of a fatty acyl chain amidated with an amino containing group^27^. Specifically, we observed an increase in the levels of short-chain *N*-acyl lipids, including acetate (C2:0) and propionate (C3:0), across different head groups (LME: all p < 0.01; **Fig. 1i**). Finally, we observed distinct developmental trajectories for the acylcarnitines based on their chain length (**Fig. 1j**). These compounds are possibly originating from host-microbial co-metabolism where microbially-produced short-chain fatty acids^35^ are attached to carnitine. Long-chain (C16-C18) acylcarnitines increased during the first month of life (LME: all p < 0.001), whereas no changes were observed for most short-chain (C3-C5) acylcarnitines (LME, p > 0.05). This is likely due to maternal transfer by breastfeeding, which was supported by the detection of these short-chain acylcarnitines in human milk.

These metabolic changes co-occurred with maturation of the infant gut microbiome. Using the phylogenetic robust center log ratio (rclr) metric phylo-RPCA^36^ to assess beta diversity, we demonstrated that age influenced infant fecal microbiota composition (PERMANOVA: F-statistic = 18.3, p < 0.001; **Fig. 2a)**. Since the difference was most pronounced between samples collected before and after one month (30 days) of age, that time point was chosen as the inflection point for subsequent categorical differential abundance analysis. Using ANCOM-BC2, we identified specific taxa driving maturation of the gut microbiome (p*_adj_* < 0.05; **Fig. 2b** for top/bottom 12 microorganisms). Early gut colonizers such as *Enterococcus^37^* decreased over time, while several *Bifidobacterium* species became more abundant after the first month of life. Using the genera identified by ANCOM-BC2 (**Fig. 2b, Extended Data Fig. 3a**), we calculated a natural log ratio of genera enriched after one month (numerator) to genera depleted before one month (denominator) (**Fig. 2c**). An approximately 3-log fold increase was observed in older compared to younger infants (Day 0 mean ratio = -1.05; Day 60 mean ratio = 2.14) with a positive trend across all major timepoints (LME: β = 0.026, p < 0.001). While Faith’s phylogenetic diversity (PD) for infant fecal samples did not change over the study period (**Extended Data Fig. 3b-c**), there appeared to be a trend of lower values for the first 60 days that then began to increase, potentially due to antibiotic exposure.

**Fig. 2:**
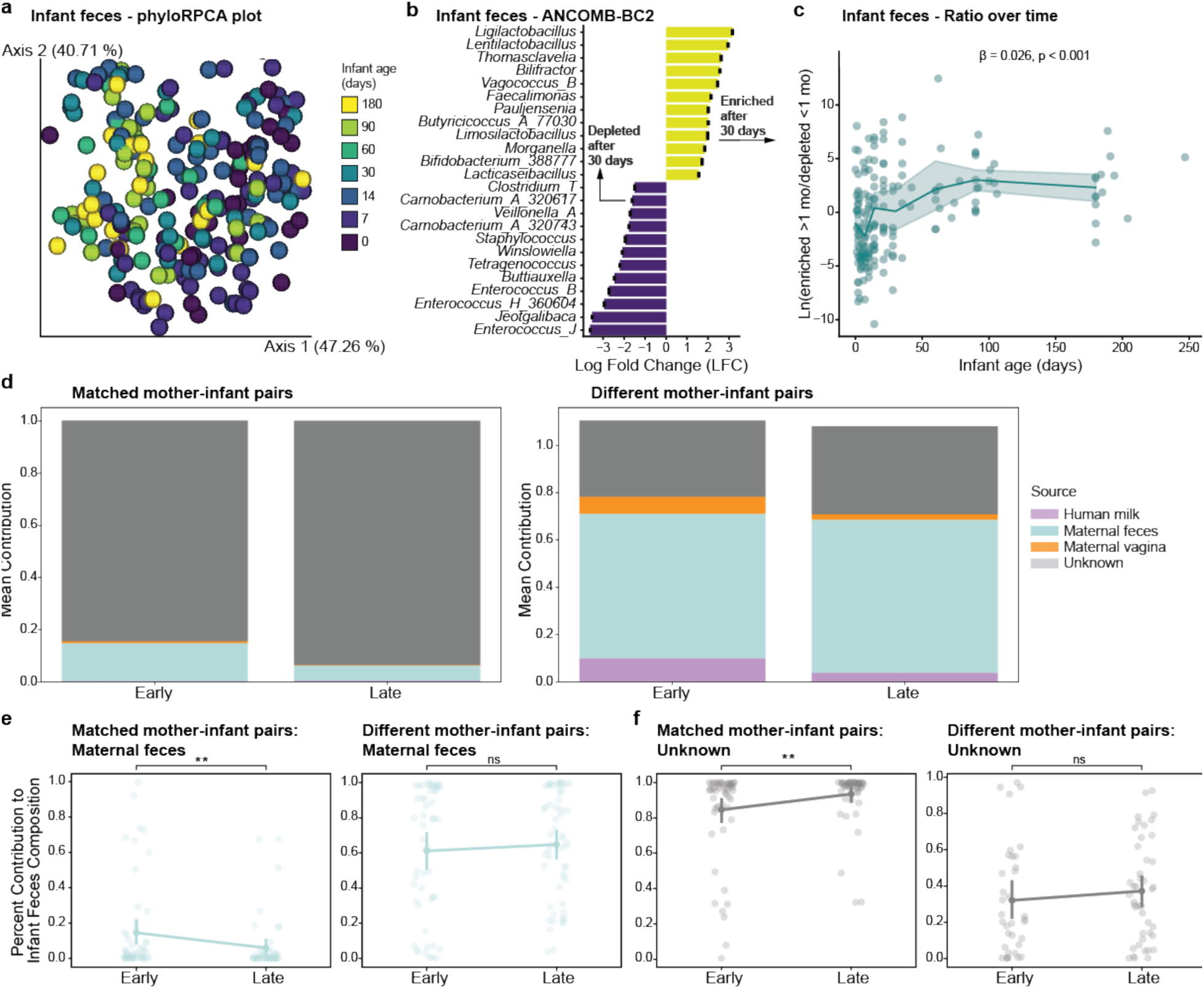
Sources driving the developing gut microbiome. **a,** phylo-RPCA EMPeror plot of fecal samples with more than 100,000 reads (n=219) colored by infant age bin. **b,** Top and bottom 12 differentially abundant infant fecal genera identified by ANCOM-BC2 (p_adj_ < 0.05, LFC > 0.5). Full plot shown in **Extended Data Fig. 3a**. Enriched (yellow) or depleted (purple) after 30 days (later timepoints) are shown, black error bars indicate standard error. **c,** Scatterplot with lineplot of the natural log-ratio of the summed counts of enriched (numerator) and depleted (denominator) fecal genera found in **b** over time (LME: β = 0.026, p < 0.001); lineplot used binned timepoints. <u>FEAST</u>: Infant fecal samples as sinks. **d,** Stacked barplot of the mean percentage contribution of each maternal body site on infant fecal samples for matched (left) or all available (right) maternal samples. **e,** Stripplot/pointplot of the FEAST estimated percentage contribution of maternal fecal samples to matched (left) or all available (right) infant fecal samples. **f,** Stripplot/pointplot of the FEAST estimated percentage contribution of unknown elements to matched (left) or all available (right) infant fecal samples. Notation: ** = p < 0.01.

Further, we observed that Faith’s PD alpha diversity decreased for infant skin samples (LME: p < 0.001; **Extended Data Fig. 3d-e**) and increased for infant oral samples (LME: p < 0.001; **Extended Data Fig. 3f-g**) over time. Maternal fecal samples also displayed significant changes over the course of the study (LME: p < 0.001), but primarily in the first weeks after giving birth (**Extended Data Fig. 3h**), likely related to postpartum antibiotic administration. As expected, there was a difference in Faith’s PD between maternal and infant fecal samples during the study (LME: p = 0.012; **Extended Data Fig. 3i**). There was no difference in the adult vaginal microbiome at the time of birth between delivery modes (**Extended Data Fig. 3j**). Similarly, there were no changes in the human milk microbiome over time (**Extended Data Fig. 3k**). All samples are compared using two beta diversity metrics, phylo-RPCA and Unweighted Unifrac, which showed differences between all body sites (PERMANOVA: p < 0.05 for all; **Extended Data Fig. 4a,b**). Finally, when looking at the phylo-RPCA beta diversity of infant samples, we see overlap between body sites at birth (Day 0) but by 6 months (Day 180) we see body sites are separated (PERMANOVA: p*_adj_* < 0.01 for all; F-statistic: feces-skin 265, skin-tongue 195, feces-tongue 7) as the microbes adapt to each niche (**Extended Data Fig. 4c,d)**.

### Maternal and Environmental Sources Seed the Developing Gut Microbiome

To explore sources of microbiota colonization, we performed source tracking using FEAST (Fast Expectation-mAximization microbial Source Tracking)^38^. When using both matched and all available maternal samples as sources, FEAST estimated that, of the available biospecimen types, maternal feces contributed the most to infant fecal microbiome development (**Fig. 2d**). Matched mother-infant samples had a much larger proportion of unknown, likely environmental, contributions (mean unknown proportions: early = 0.85, late = 0.94), than when using all available adult samples as possible sources of microbes (mean unknown proportions: early = 0.32, late = 0.37). This was further supported by the fact that there were no differences in phylo-RPCA beta diversity differences within and between related mother-infant pairs (**Extended Data Fig 4e,f**). There was no significant difference in the proportion of the infant gut microbiome attributed to matched or all available maternal vaginal or human milk samples when comparing early and later timepoints (**Extended Data Fig. 4g,h**), indicating that bacteria seeded at birth did appear to persist, although there was individual infant variation. There was also a decrease in the proportion of the infant gut microbiome attributed to matched maternal fecal samples between early and late timepoints (2MWW, p = 0.0068), but no significant differences in proportions over time when considering all available maternal fecal samples as possible sources (**Fig. 2e**). When non-matched maternal fecal samples were considered, they appeared to contribute proportionally more to the infant gut microbiome (approximately 60%) compared to when only matched maternal fecal samples were examined (approximately 10%) (**Fig. 2e**). This suggests that environmental sources, including other household members and non-maternal caretakers^39,40^, play a larger role in the ongoing development of the infant microbiome than previously understood. There was also a significant increase in the contributions of unknown sources over time on the infant fecal microbiome with matched samples (2MWW, p = 0.0042), but not when considering all available sources (**Fig. 2f**). In general, the infant gut microbiome development trajectory appears to mature towards an adult-like state, rather than converging towards the mother’s specific microbiome as previously suggested^6,41,42^. Together, this indicates that while maternal seeding is important, a large proportion of the infant gut microbiome is derived from increasing amounts of environmental sources over time, at least in this Bangladeshi cohort.

### Cesarean Delivery is Associated with Higher Levels of Microbially Derived Metabolites in Early Infancy

While infant age was the primary variable shaping the fecal metabolome, PERMANOVA also identified other significant factors, including delivery mode (**Fig. 3a**). The model output for all tested variables is provided in **Supplementary Table 2**. The natural log ratios of features associated with C-section (numerator) and vaginal birth (denominator) identified by PLS-DA (CER = 0.40; **Extended Data Fig. 5a**) separated groups over time (LME: β = -0.71, p = 2.4×10^-5^; **Fig. 3b**), with the largest difference observed during the first month of life. Infants born via C-section were enriched in amino acids, dipeptides, acetylcarnitine (C2:0), and N-acetylneuraminic acid (Neu5Ac). Neu5Ac is the primary sialic acid component found within sialylated HMOs in human milk^15^, with roles in host-microbe interactions^43^ and neurodevelopment^44,45^. Conversely, vaginally delivered infants showed enrichment of several microbially derived bile acids and saccharides (**Supplementary Table 1**). Furthermore, different *N*-acyl lipids were enriched in C-section and vaginally born infants, varying in head group and chain length (**Supplementary Table 1**). Surprisingly, the effect of delivery mode on infant gut microbiome composition was less pronounced, possibly due to the fact that all mothers received postpartum antibiotics, only reaching statistical significance at Day 7 (2MWW-HB, p = 0.006; **Extended Data Fig. 5b**), but not over the entire study period (LME: p = 0.118). However, we did see a statistically significant effect of delivery mode on Faith’s phylogenetic alpha diversity over time (LME: β = 11.8, p = 0.029).

**Fig. 3:**
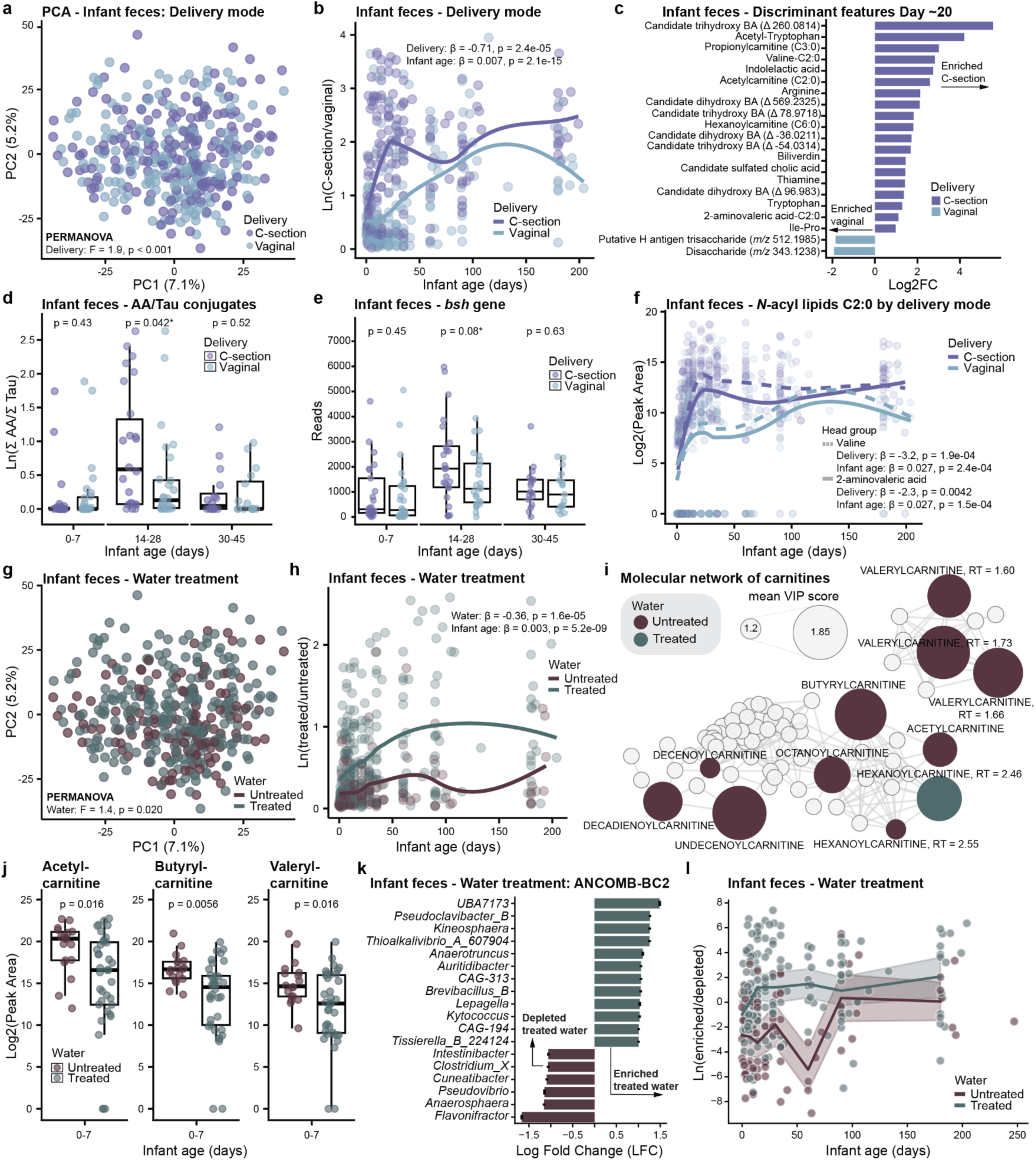
Delivery mode and water treatment are associated with infant fecal metabolic trajectories in early-life. **a,** PCA of infant fecal samples showing separation by delivery mode (PERMANOVA: p < 0.001). **b**, Longitudinal natural log ratios of the summed peak areas of the top 100 discriminating features for delivery mode in infant feces. LME was used to determine statistical significance between groups. Scatter plot smoothing (‘LOESS’) and removal of three outliers for visualization. **c**, Barplot of metabolic features significantly differing between C-section and vaginal birth (p*_adj_* < 0.10) in infant feces collected during days 10-30 after birth. For each infant, the sample with the highest log ratio within this interval was selected. **d**, Boxplot of the natural log ratios of amino acid to taurine conjugated bile acids by delivery mode at selected timepoints (Wilcoxon rank-sum test, day 14-28: p = 0.042, p*_adj_* = 0.13). **e**, Boxplot of *bsh* gene reads in infant feces by delivery mode (Wilcoxon rank-sum test, day 14-28: p = 0.07, p*_adj_* = 0.08). **f**, Trajectories for valine-C2:0 and 2-aminovaleric acid-C2:0 by delivery mode in infant feces, with smoothing for visualization. The difference between C-section and vaginal birth was assessed for significance using LME. **g**, PCA of infant fecal samples showing separation by household drinking water treatment (PERMANOVA: p = 0.020). **h**, Longitudinal natural log ratios of the summed peak areas of the top 100 discriminating features for treated vs. untreated drinking water. Smoothing added for visualization, and statistical significant difference between groups was assessed using LME. **i,** Molecular network of acylcarnitines associated with household drinking water treatment (untreated=brown, treated=dark green). Node size represents mean VIP score from three different PLS-DA models accounting for different purification methods, with larger nodes indicating higher VIP score. Only acylcarnitines that were significant in all three models (VIP > 1) and with the same directionality were included in the network. **j,** Boxplot showing differences in the natural log peak areas of various acylcarnitines enriched in the untreated water group at day 0-7. Statistical significance was determined by the Wilcoxon rank-sum test with BH adjusted p values. **k,** Differentially abundant infant fecal genera by water treatment identified by ANCOMB-BC2 (q < 0.05, log fold change (LFC) > 1). Enriched (grey) or depleted (brown) in treated water are shown, black errors indicate standard error. **l,** Scatterplot/lineplot of the natural log-ratio of the summed counts of enriched (numerator) and depleted (denominator) fecal genera found in **k** (LME: -0.058, p = 0.13); lineplot used binned timepoints. For **b, f,** and **h,** β and p values were derived from LME models with subject as random effect. All boxplots show the first (lower), median, and third (upper) quartiles, with whiskers 1.5 times the interquartile range. For boxplots, repeated measures were avoided by retaining only the highest value per individual within each time window.

To further explore the divergence in metabolic trajectories based on delivery mode, we focused on days 10-30 after birth, the earliest point of group separation. Most discriminant features during this window were enriched in infants delivered via C-section and included bile acids, short-chain acylcarnitines, *N*-acyl lipids, amino acids, tryptophan derivatives, dipeptides, and lipid derivatives (**Fig. 3c**). Compounds such as indolelactic acid and dipeptides are known metabolic products of the gut microbiome^46,47^, often generated through microbial digestion of proteins into peptides and amino acids. In contrast, features enriched in vaginally delivered infants included milk-associated glycans (**Fig. 3c**). Given the observed differences in several bile acids, we further investigated the ratio of microbiome-derived amino acid to host-derived taurine conjugates and the levels of microbial *bsh* gene in infant stool. Interestingly, we observed a trend where both this ratio and *bsh* gene levels were transiently elevated in C-section compared to vaginally born infants between days 14-28, though these differences did not reach statistical significance after FDR correction (Wilcoxon rank-sum test, p_adj_ = 0.13 and p_adj_ = 0.08, respectively; **Fig. 3d,e**). This may indicate a temporary enrichment of microorganisms with more *bsh* genes in C-section born infants the first weeks of life, however, this needs further investigation in a larger cohort. Further, we observed transiently elevated levels of valine-C2:0 (LME: β = -3.2, p = 1.9×10^-4^) and 2-aminovaleric acid-C2:0 (LME: β = -2.3, p = 0.0042) in C-section vs. vaginally born infants (**Fig. 3f**). A similar pattern was observed for acetylcarnitine (C2:0), propionylcarnitine (C3:0), and hexanoylcarnitine (C6:0), with higher levels between days 14-28 (Wilcoxon rank-sum with BH correction, all p < 0.05; **Extended Data Fig. 5c**). Although we observed only minor differences in the gut microbiome composition based on delivery mode, its metabolic output may differ, as suggested by the distinct profiles obtained for the microbially derived metabolites, including those involved in immune regulation pathways, such as bile acids^30^ and *N*-acyl lipids^27^. This may contribute to differences in early-life immune maturation observed between delivery modes^8,48^. Indeed, early-life changes in bile amidates have recently been found to be predictive of islet autoimmune dysregulation in diabetic onset post-birth while *N*-acyl lipids controlled T-cell differentiation^49^.

### Household Drinking Water Treatment is Associated with Altered Carnitine Metabolism

Consistent with the source tracking analyses showing a significant environmental contribution to the infant gut microbiome, both socioeconomic status (asset index, PERMANOVA: F-statistic = 1.5, p = 0.006; **Extended Data Fig. 5d-f**) and reported household drinking water treatment practices (PERMANOVA: F-statistic = 1.4, p = 0.020; **Fig. 3g**) were associated with different infant fecal metabolic profiles. We further explored the relationship to drinking water treatment, viewing it as an indirect proxy for household hygiene and sanitation practices, given previously reported association with the gut microbiome^50,51^. The natural log ratios of features associated with treated water (numerator) and untreated water (denominator) identified by PLS-DA (CER = 0.35; **Extended Data Fig. 5g**) clearly separated groups throughout the study period (LME: β = -0.36, p = 1.6×10^-5^; **Fig. 3h**). To account for different purification methods (e.g. boiling vs. filtering), we built three separate PLS-DA models (**Extended Data Fig. 5h-i**). Despite that the overall model performances were modest (CER < 0.45), intersecting features with a VIP > 1 and same directionality revealed 255 common metabolic features (**Supplementary Table 3**). The most prominent finding was a consistent enrichment of multiple acylcarnitines in the untreated water group (**Fig. 3i**), compounds central to fatty acid metabolism, host mitochondrial function, and energy metabolism^52^. Interestingly, the difference in carnitine metabolism was most pronounced the first weeks of life (**Fig 3j**). Given that acylcarnitines are present in human milk, with higher levels of acetylcarnitine (C2:0) and propionylcarnitine (C3:0) in colostrum as shown previously, we investigated if water treatment also influenced the milk metabolome. Although not significant in the PERMANOVA (F = 1.2, p = 0.08), a PLS-DA model (CER = 0.32, **Extended Data Fig. 5j**) identified several acylcarnitines as discriminant features between treated and untreated water (**Supplementary Table 1**). Notably, we found an opposite pattern than we observed for infant feces, with acylcarnitines being depleted in human milk in the untreated water group (exemplified for acetylcarnitine (p < 0.01) and propionylcarnitine (p < 0.01) in **Extended Data Fig. 5k**). These observations raise an intriguing hypothesis: that water treatment methods may have a direct impact on metabolic profiles. While this warrants further investigation, it is possible that water treatment serves as a proxy for broader socioeconomic differences, particularly given similar patterns observed with the asset index used as a socioeconomic proxy.

Water treatment was weakly associated with overall gut microbiota composition, but it was linked to changes in specific genera, particularly in early life (**Fig 3k,l**). Infants in the untreated water group had enriched amounts of genus *Flavonifractor* (some members formerly *Clostridium)*, known for its ability to metabolize flavonoid polyphenols^53^, and a current *Clostridium* genus, which is known to play a role in the metabolism of carnitines and its related metabolites^54^. Consistent with this, a search of the spectra from discriminant acylcarnitines using microbeMASST^55^, identified matches to bacteria including *Clostridium* (exemplified for butyrylcarnitine in **Extended Data Fig. 5l**). The elevated levels of carnitines in infant feces in the untreated water group may therefore be attributed to a microbiome enriched in species able to produce them, such as *Clostridium* species. However, it is also possible that higher availability of acylcarnitines in untreated water promotes the growth of carnitine-metabolizing bacteria. We also found that infants in the untreated water group had significantly lower phylogenetic alpha diversity in the first 30 days of life (2MWW-HB, p = 0.0072; **Extended Data Fig. 5m**), suggesting a potential negative impact on the establishment of a diverse microbiome, which may have implications for long-term health given that lower microbial diversity has been associated with various diseases^56,57^.

### Maternal Secretor Status Influences Infant Fecal Metabolic Programming

We also investigated the influence of maternal secretor status on both the human milk and the infant fecal metabolome. As expected and confirmed by targeted quantification using high-performance liquid chromatography (HPLC) with fluorescent derivatization, there was a clear difference in HMO milk profiles between secretors and non-secretors (**Extended Data Fig. 6a-b**), consistent with previous findings^58–60^. Surprisingly, unsupervised dimensionality reduction of the milk metabolome did not reach statistical significance for maternal secretor status, but a trend was observed (PERMANOVA: R^2^ = 0.01, F-statistic = 1.2, p = 0.067). Nevertheless, by plotting the natural log ratio of discriminant features associated with secretor vs. non-secretor identified by PLS-DA (CER = 0.36; **Extended Data Fig. 6c**) we observed a consistent separation over time (LME: β = 1.0, p = 5.1×10^-13^; **Fig. 4a**). The difference was largely driven by features putatively classified as saccharides by SIRIUS^61,62^. By molecular networking and manual inspection of MS/MS spectra, we identified several putative HMOs, including 2’FL (*m/z* 471.1709) and difucosyllactose (DFLac) (*m/z* 657.2211), another secretor-associated HMO^63,64^. As expected, these were detected almost exclusively in milk samples from secretors (**Fig. 4b**). While saccharides such as 2’FL cannot be unambiguously differentiated from their structural isomers by MS/MS alone, their exclusive presence in secretors supports their annotation as α1-2-fucosylated HMOs. Additionally, the observed patterns were consistent with the targeted quantification of 2’FL and DFLac (**Fig. 4b**), and in secretor milk, the targeted data correlated with the untargeted data for these compounds (Pearson correlation: 2’FL: r = 0.65, p = 1.8×10^-5^ and DFLac: r = 0.84, p = 4.2×10^-10^; **Extended Data Fig. 6d**).

**Fig. 4:**
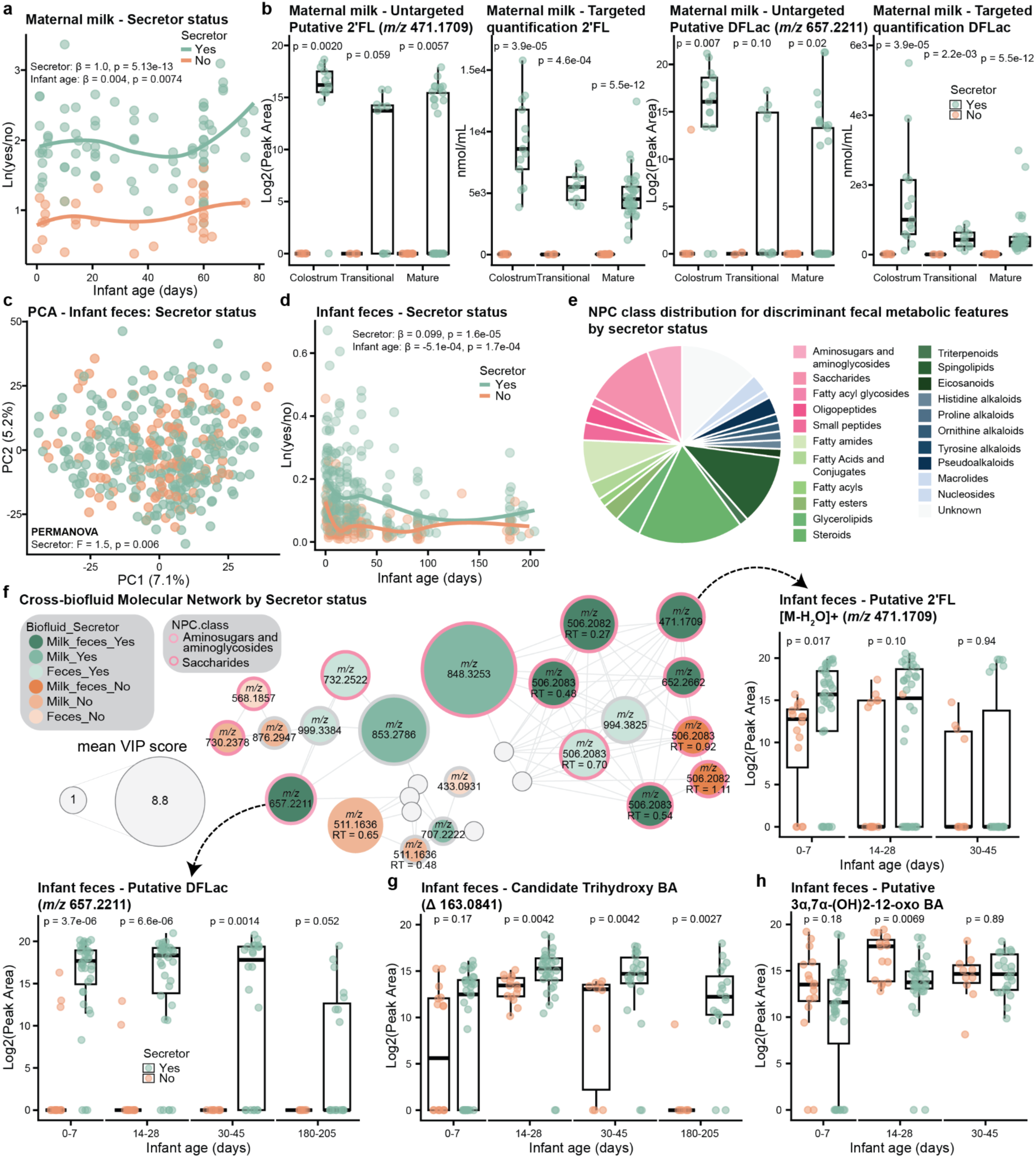
Maternal secretor status shapes HMO profiles in human milk and infant feces. **a,** Longitudinal natural log ratios of the summed peak areas of the top 70 discriminating features for maternal secretor status in human milk. Statistical significant difference between groups was assessed with LME. Scatter plot smoothing (‘LOESS’) added for visualization. **b,** Boxplot of metabolic feature (*m/z* 471.1709) predicted to be 2’FL, targeted quantification of 2’FL, metabolic feature (*m/z* 657.2211) predicted to be DFLac, and targeted quantification of DFLac, respectively, grouped by secretor status (Wilcoxon rank-sum test, with BH adjusted p values). For subjects with multiple samples within a stage, only the sample with the highest 2′FL concentration was retained. **c,** PCA of infant fecal samples showing separation by maternal secretor status (PERMANOVA: p = 0.006). **d,** Longitudinal natural log ratios of the summed peak areas of the top 100 discriminating features for maternal secretor status in infant feces, with the significance of the difference assessed by LME and smoothing for visualization. **e,** Piechart showing the distribution of NPC predicted chemical classes or pathways (probability > 0.7) for the discriminant infant fecal features by maternal secretor status. Only features with VIP > 2.5 were included. **f,** Cross-biofluid molecular network of amino sugars, aminoglycosides, and saccharides, as predicted by CANOPUS, colored by biofluid and secretor status. Node size represents the mean VIP score of significant features across the two PLS-DA models for secretor status in human milk and infant feces, with larger nodes indicating higher VIP score. Selected features enriched in both maternal milk and infant feces are shown in boxplots. **g,** Boxplot of the natural log-transformed peak areas of a candidate trihydroxy bile acid (Δ 163.0841) grouped by secretor status (Wilcoxon rank-sum test with BH adjusted p values). **h,** Boxplot of the natural log-transformed peak areas of putative 3α,7α-(OH)_2_-12-oxo based on secretor status at select timepoints (Wilcoxon rank-sum test with BH adjusted p values). All boxplots show the first (lower), median, and third (upper) quartiles, with whiskers 1.5 times the interquartile range. For boxplots in **f, g,** and **h**, repeated measures were avoided by retaining only the highest value per individual within each time window. For **a** and **d,** β and p values were derived from LME models with subject as random effect.

The difference in human milk composition between secretors and non-secretors drove a corresponding association with the infant fecal metabolome (PERMANOVA: F-statistic = 1.5, p = 0.006; **Fig. 4c**). The natural log ratio of discriminant features associated with secretor status identified by PLS-DA (CER = 0.34; **Extended Data Fig. 6e**), revealed that infants of secretor mothers had distinct fecal metabolic profiles from those of non-secretor mothers in the first months of life (LME: β = 0.099, p = 1.6×10^-5^; **Fig. 4d**). The metabolic difference diminished around 3-4 months of life, likely reflecting reduced breastfeeding, as less than 45% of infants were still exclusively breastfed at that time. Key discriminant features included not only saccharides (11.4%), but also steroids (17.1%), sphingolipids (11.4%), and aminosugars/aminoglycosides (5.7%) (putatively classified by CANOPUS; **Fig. 4e**), suggesting a broad effect of human milk composition on gut metabolic function. For example, dipeptides and microbial bile amidates, including glutamate-trihydroxy and glutamine-trihydroxy, were enriched in the feces of infants born to secretor mothers (**Supplementary Table 1**). N-acetyl-D-glucosamine (GlcNAc), an amino sugar known to be an important energy source for gut bacteria^65^, were enriched in infants of non-secretors, supporting previous findings of lower GlcNAc in infant feces from secretor mothers^66^. Furthermore, acylcarnitines, histamine-trihydroxy bile acid, several short-chain *N*-acyl lipids, and polyamine derivatives (e.g. spectral matches to acetylputrescine-C2:0, cadaverine-C3:0, acetyl-palmitoyl-putrescine, and diacetylspermidine) were also enriched in non-secretors, whereas *N*-acyl lipids enriched in secretors were predominantly longer-chain compounds (C16-C18) (**Supplementary Table 1**). Although the source tracking analysis showed that direct microbial seeding from maternal milk is limited, these findings highlight the indirect effect of HMOs in shaping gut metabolic programming^67^. A cross-biofluid molecular network demonstrated transfer of specific HMOs from human milk to infant feces, showing that HMOs abundant in secretor milk, including putative 2’FL and DFLac, were also enriched in the feces of respective infants (**Fig. 4f**). This suggests that maternal secretor status directly shapes the infant fecal metabolome through different composition of the milk itself, rather than simply differences in breastfeeding patterns.

Interestingly, secretor status also influenced bile acid metabolism. Notably, a candidate trihydroxy bile acid (Δ 163.0841) was consistently elevated in infants of secretor mothers throughout the study period (Wilcoxon rank-sum test: all p *<* 0.01), except during the first postnatal week (**Fig. 4g**). We identified it as a bile acid given the two diagnostic ions, *m/z* 319.24 and 337.25, in the MS/MS spectrum^30^, with delta mass molecular formula C_6_H_13_NO_4_, as predicted by SIRIUS. We do not yet know the structure of the bile acid steroid core or the modification as it is a spectral match to the candidate bile acid library^30^ and therefore it is referred to as a candidate bile acid. Conversely, the microbial-derived 3α,7α-(OH)_2_-12-oxo bile acid, characterized by a ketone at the C12-position and formed through oxidation by bacterial 12α-hydroxysteroid dehydrogenases^68^, exhibited a transient difference, with higher levels in infant feces of non-secretors between days 14 and 28 postpartum (Wilcoxon rank-sum test, p = 0.007; **Fig. 4h**). Secretor status did not influence the infant fecal microbiome alpha diversity (Faith’s PD, LME: β = 8.1, p = 0.13) nor the fecal log ratio from **Fig. 2c** (LME: β = 0.001, p = 0.94; **Extended Data Fig. 6f**).

### Identification of Microbial Drivers of Early-life Metabolic Programming through Multi-omics Analysis

Finally, to identify candidate bacteria associated with the production of metabolites of interest, we performed a multi-omics analysis using joint-RPCA^36^. The joint analysis was restricted to infant fecal samples collected between days 10-30 after birth, roughly aligning with the earliest time window showing group separation by the variables investigated. We focused on microbial-modified bile acids, *N*-acyl lipids, acylcarnitines, and other key differential metabolites identified in earlier analyses. Microbiome features were selected based on the overlap between three analyses (ANCOM-BC2, joint-RPCA, and TEMPTED). The joint-RPCA found delivery mode as a significant factor shaping combined multi-omics analysis (PERMANOVA: delivery mode, p = 0.01, F-statistic = 3.98; **Extended Data Fig. 7a**). Further, there were strong positive or negative correlation patterns with key features of interest **(Fig. 5a**; complete heatmap in **Extended Data Fig. 7b**). Notably, several compounds associated with fatty acid metabolism, including microbial bile amidates, 2-aminovaleric acid-C2:0, valine-C2:0, and several acylcarnitines negatively co-occurred with OGUs belonging to *Escherichia* and *Bifidobacterium* genera. This may indicate that these molecules can be used by these genera for growth or other metabolic functions. Consistent with other reports^16,17^, we observed a positive correlation between *Bifidobacterium* and HMOs. Preliminary evidence suggests that *Escherichia* and *Bifidobacterium* can act cooperatively to digest HMOs^69^. Furthermore, since *Bifidobacterium* spp. are critical members of the early-life gut microbiome^70,71^, and are involved in HMOs utilization, it was interesting to note that they also displayed strong positive co-occurrences with biliverdin, several acylcarnitines, 3α,7α-(OH)_2_-12-oxo bile acid, and at least one sulfated trihydroxy bile acid. This suggested that *Bifidobacterium* spp. may produce or co-metabolize these compounds. To assess this further, we searched the MS/MS spectra of these metabolites with microbeMASST^55^. Overall, we obtained a match for 68% of the searched spectra, suggesting that these bacteria could actually produce the molecules of interest. Importantly, 53% of them had matches to monocultures from the same genera to which they were associated in the heatmap (**Fig. 5a**). For example, acetylcarnitine (C2:0) was frequently observed in cultures of *Bifidobacterium* (**Extended Data Fig. 7c**).

**Fig. 5:**
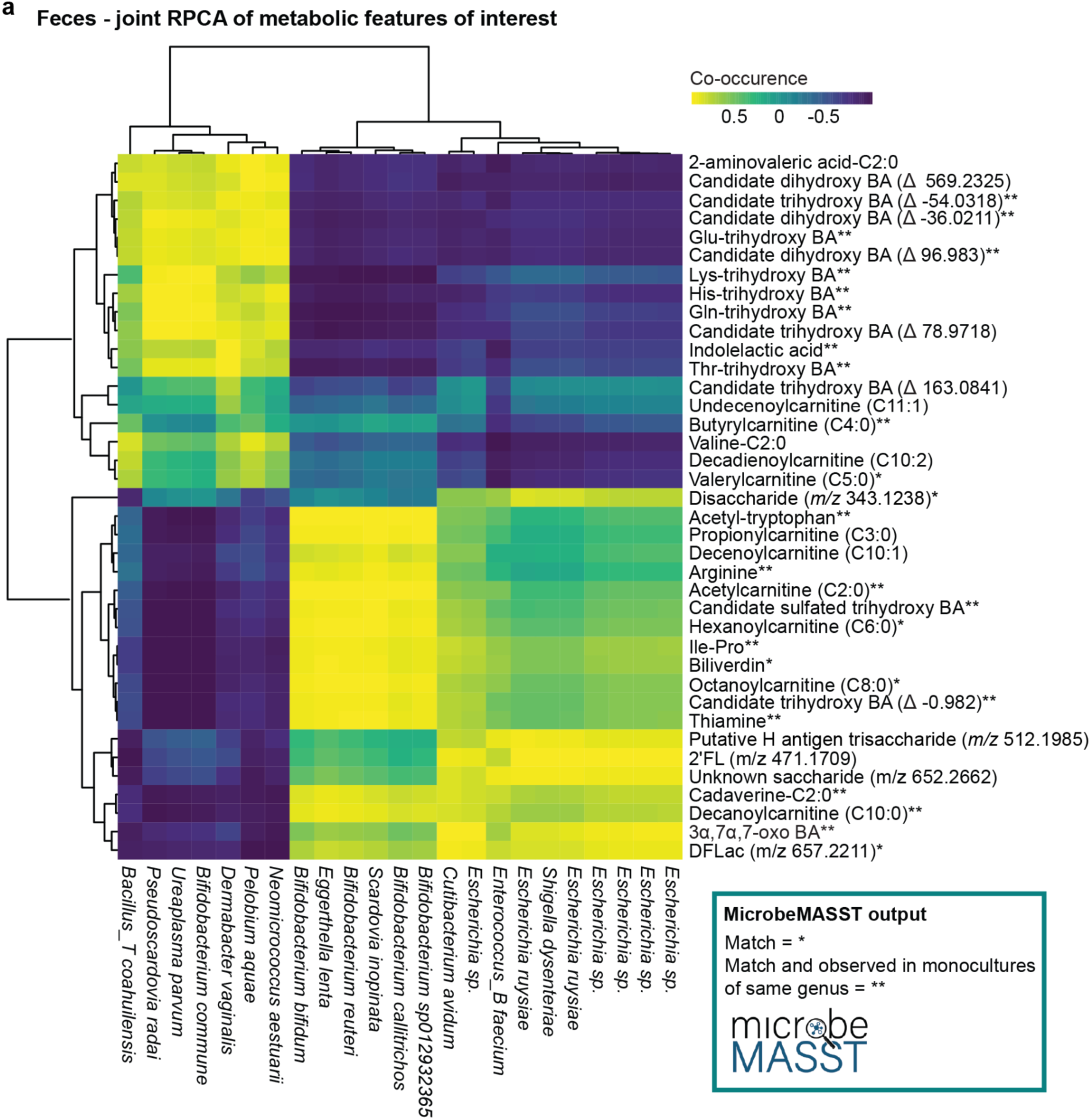
Multi-omics integration identifies microbial mediators of metabolic trajectories. **a,** Heatmap of correlating metabolites and microbial species from infant fecal samples collected at day ∼30 identified by joint-RPCA. Metabolic features shown were selected based on previous analyses. Asterisk indicate MS/MS spectral match against microbeMASST, with (*) indicating a general match and (**) indicating a match to a monoculture of the same genus as taxa shown in the heatmap. Microbiome features shown were selected based on the overlap between 3 previous analyses, including needing to belong to the differentially abundant genera found in the ANCOM-BC2 analysis from Figure 2b, the top and bottom 20 most differentially ranked features for joint-RPCA Axis 1, and the top and bottom 10% differentially ranked features from TEMPTED Axis 1 (overlapping features=23).

## DISCUSSION

This study, including over 1,600 metagenomics samples and more than 500 untargeted metabolomics profiles across multiple body sites, offers unique and comprehensive insights into infant metabolic programming during early life in a cohort from Bangladesh. Through frequent sampling over the first six months of life, we profiled the metabolome and microbiome of 55 mother-infant dyads. Leveraging recently developed MS/MS spectral libraries^27,30–32^, we describe the fecal developmental trajectories of key signalling molecules, including bile acids, *N*-acyl lipids, and acylcarnitines. Through multi-omics integration we also reveal specific microbial-metabolite relationships, emphasizing the role of the gut microbiome as an important mediator of early developmental metabolic trajectories.

We show that the functional capability of the gut microbiome matured rapidly, demonstrated by an early rise in microbially derived bile acids and short-chain *N*-acyl lipids in the first weeks after birth. This reflects the developing microbial proficiency in performing modifications of these compounds. The decline in host-derived taurine and glycine conjugates alongside the rise in microbial bile acids aligns with previous findings^33,72^, suggesting a shift toward a microbiome capable of more complex bile acid transformations, such as deconjugation and amidation^30^. This was further supported by increased levels of the *bsh* genes with infant age, reflecting colonization of bsh-expressing bacteria as the infant grows^33^. Understanding the developmental trajectories of these compounds is crucial as bile acids not only interact with nuclear receptors such as farnesoid X receptor (FXR)^33,73^ and pregnane X receptor (PXR)^31,74^, but also modulate immune processes, including T-cell differentiation and dendritic cell signaling^31,33,74^, thereby shaping and programming infant immune development.

Further, we demonstrate that the trajectories of these metabolites are sensitive to factors like delivery mode and maternal secretor status. Despite only minor differences in infant gut microbiota composition, infants born via C-section had unique metabolic profiles, with transiently higher levels of bile amidates and *N*-acyl lipids, as well as short-chain acylcarnitines, during the first month of life. Beyond their critical role in energy metabolism and mitochondrial homeostasis^52,75^, acylcarnitines have been associated with rapid infant growth^70^, and they are also involved in immune homeostasis^52^ by modulating inflammatory signalling pathways, such as NF-κB and JNK^76,77^. Second, we found that maternal secretor status also influenced the bile acid pool, with levels of a candidate trihydroxy bile acid (Δ 163.0841) and 3α,7α-(OH)_2_-12-oxo bile acid differing between infants of secretor and non-secretor mothers. This suggests that the maternal HMO profiles, through the infant gut microbiome modulation^17,67,78^, may shape early-life bile acid metabolism.

Source tracking analysis revealed that, while the maternal fecal microbiome is the main contributor to the infant fecal microbiome, environmental sources also play a major role in shaping its composition. Environmental exposures such as drinking water quality can differ substantially between LMIC and high-income settings based, for example, on contamination with diarrheagenic^79^ or antibiotic-resistant pathogens^80^. Exogenous elements, including xenobiotic trace elements in water, have been linked to changes in the diversity and composition of the gut microbiome in both adults^50,51^ and infants^81^. Here, we show that differences in the infant fecal metabolome and microbiome are associated with reported household-level drinking water treatment practices, specifically in relation to samples collected during the first month of life. Given that infants usually do not directly consume water at this age, this variable should be interpreted as a proxy for household hygiene and sanitation related practices. As it is also likely associated with socioeconomic status and other unmeasured factors, this finding may reflect a broader constellation of hygiene practices, infrastructure, and living conditions, rather than water treatment *per se*. Notably, infants in households using untreated water presented higher fecal levels of acylcarnitines, despite being depleted in the milk of their respective mothers. Accumulation of long-chain carnitines in the feces has been previously associated with intestinal inflammation in the pediatric context and growth promotion of *Enterobacteriaceae* species and other pathobionts^82^. Supporting this, we found *Clostridium* species, also known to metabolize acylcarnitines^54^, enriched in infants from households that did not report using methods to treat water. Elevated levels of acylcarnitines have been linked with negative health outcomes^83–85^, but long-term health implications remain to be investigated. Another exposure that can differ substantially from high-income settings is the widespread use of antibiotics in LMICs^21–23^. This creates a different baseline for microbial development than in a Western setting, where treatment with antibiotics is less frequent^86^, potentially explaining the observed differences in alpha diversity for infant gut microbiota^41^.

In conclusion, this longitudinal multi-omics study emphasizes the influence of delivery mode, maternal secretor status, and environmental factors like water treatment in shaping the early infant microbial and metabolic profiles in a LMIC setting. This study has a few limitations, including a relatively small sample size for the dyads (n=55), COVID-19 pandemic may have impacted household habits during the study, reliance on self-reported data for variables like water treatment, widespread antibiotic usage, the lack of inclusion of seasonality in the analysis, and metabolite annotations based on MS/MS spectral matches with the GNPS spectral libraries (level 2-3). Additionally, the water source was not considered, but most households in Dhaka used piped city water or deep tube wells^87^. Despite these limitations, we show that the infant gut microbiome, a dynamic ecosystem, requires a multi-omics approach to better understand the rapidly changing metabolic mechanisms of early life that can ultimately impact the future health and development of individuals. Follow-up studies using long-read sequencing, which provides better strain-level resolution^88^, in larger cohorts will be essential for determining the extent and nature of the observed responses to both strain transmission as well as water quality.

## METHODS

### Ethics statement

Ethics approval for the SEPSiS (Synbiotics for the Early Prevention of Severe Infections in Infants) Observational Cohort Study, from which the biological specimens and data used in this study were derived, was obtained from the Research Ethics Board of The Hospital for Sick Children (No.: 1000063899) and the International Center for Diarrhoeal Disease Research, Bangladesh (icddr,b; PR-19045). In addition, this project has been reviewed by the Director of the UCSD HRPP, IRB Chair, or IRB Chair’s designee and is exempt from Institutional Review Board requirements under categories 45 CFR 46.104(d), category 4: Secondary research for which consent is not required: Secondary research uses of identifiable private information or identifiable biospecimens; ii) Information, which may include information about biospecimens, is recorded by the investigator in such a manner that the identity of the human subjects cannot readily be ascertained directly or through identifiers linked to the subjects, the investigator does not contact the subjects, and the investigator will not re-identify subjects.

### Study design and recruitment

Mother-infant pairs were screened at two government health care facilities offering delivery services in Dhaka: Maternal and Child Health Training Institute (MCHTI) and Mohammadpur Fertility Services and Training Centre (MFSTC). Enrollment began in May 2021 and the last study visit was completed in April 2022. Eligible participants were offered enrollment into ‘Schedule A’ (n=75) of the SEPSiS Observational Study, which was designed for metagenomics and metabolomics analyses (**Extended Data Fig. 1a**). Infants were eligible for inclusion if: informed consent was received by parent or guardian, they were between 0-1 days old at screening, delivered at one of the study hospitals, orally feeding at the time of screening and planned to reside within the study catchment area until 60 days of age.

Exclusion criteria were: birth weight <1500 g, major surgery required within the first week of life, and/or major congenital anomaly of the gastrointestinal tract, maternal human immunodeficiency virus (HIV) infection and/or history of mother ever receiving antiretroviral drug(s) for presumed HIV infection or administration/prescription of parenteral antibiotics or any prenatal or postpartum use of non-dietary probiotic supplement, infants that required mechanical ventilation and/or cardiac support, infants actively enrolled in clinical trials that included the use of probiotics or prebiotics, infants residing in the same household as another infant previously enrolled in the study, or any study within the research platform, who is currently <60 days of age. Field activities were overseen by the International Centre for Diarrheal Disease Research, Bangladesh (icddr,b). Participants were followed from enrollment to 6 months of age.

A total of 55 mother-infant pairs were selected from the 75 infants enrolled in the SEPSiS Schedule A arm and included in this study. In addition to the general inclusion and exclusion criteria described in the study design section, infants were specifically selected to contribute samples to this study if the following additional criteria were met: if day 0 or 1, 60, 90 and 180 stool samples were collected and available, at least 11 routine clinical assessments were completed, 7 or more stool samples were collected, a minimum set of data collection modules relevant for the study analyses were completed (e.g., feeding practice, birthweight, gestational age information). Enrollment was stratified by delivery mode to ensure a similar proportion of infants delivered vaginally and via C-section.

### Sample collection

Samples were collected longitudinally from various body sites by trained study personnel during routine visits in the hospitals until discharge or in the participant’s home after discharge. Eleven fecal samples were scheduled for collection from mothers and infants on days 0 or 1, 3 or 4, 6 or 7, 10, 14, 21, 28, 35, 60, 90, and 180, with allowances for rescheduling up until the next scheduled visit. Infant oral (tongue) and skin swabs (elbow) were scheduled at 8 different timepoints (day 0 or 1, 6 or 7, 14, 21, 35, 60, 90, 180), with allowances for rescheduling up until the next scheduled visit. Mothers provided a vaginal swab at enrollment (day 0/1). Maternal milk was collected at enrollment (0/1), day 60, and 2 other randomly determined timepoints up to day 60 of the infant’s age. Infant blood samples were collected at day 1 of age only.

Infant stool samples were collected by trained study personnel, aliquoted and stored in the vapour phase of liquid nitrogen within 20 minutes of defecation. Maternal fecal samples were collected and placed in a cold box within 20 minutes of defecation and then homogenized, aliquoted and frozen by a study worker within 6 hours of defecation. Vaginal swabs were self-collected by mothers at each study hospital after delivery following the instructions provided by study nurses and physicians. Mothers hand-expressed human milk in sterile containers which were aliquoted and stored on ice. Infant oral (from the tongue dorsum) and skin swabs (from the antecubital fossa, on the inner left or right elbow) were collected aseptically by trained study workers and placed in sterile containers. One routine venous blood sample was collected from infants by a trained phlebotomist, study nurse or medical officer. All specimens were transported to the International Centre for Diarrheal Disease Research, Bangladesh (icddr,b) and stored in -70°C until they were transported on dry ice from Dhaka to University of California San Diego (UCSD) for metagenomic, metabolomics, and HMO laboratory analysis.

### Main study variables and definitions

To assess contribution of various maternal and infant characteristics to the variation in the infant fecal metabolome we performed a permutational multivariate analysis of variance (PERMANOVA)^89^. Infant age was the primary variable of interest to describe developmental trajectories of key signalling molecules. In addition to infant age, the PERMANOVA model included variables selected for their known influence on the early life gut microbiome and metabolome: delivery mode (categorical)^6,8^, maternal secretor status (categorical)^17,67^, gestational age (continuous)^90^, feeding patterns (categorical)^91,92^, infant sex (categorical)^93^, birthweight (continuous)^93^, antibiotic usage by mother and by infant (categorical)^7,20^, socioeconomic status (categorical)^94,95^, and household water treatment (categorical)^50,51^. Based on the PERMANOVA output, three significant covariates were prioritized for in depth-analysis: delivery mode, maternal secretor status, and household water treatment.

- **Delivery mode:** categorized as vaginal or C-section based on clinical records.
- **Maternal secretor status:** categorized as secretor or non-secretor based on presence or near-absence (<100 nmol/mL) of HMO 2’-fucosyllactose.
- **Household drinking water treatment**: this binary variable was defined by self-reports collected at baseline. Households were categorized as ‘treated’ if they reported purifying their drinking water (either using boiling and/or filtering) or ‘untreated’ if they reported no treatment of their drinking water.

Additionally, we briefly assessed the impact of socioeconomic status on the infant fecal metabolome using an asset index. The asset index and quintiles were generated using PCA for all participants enrolled in the SEPSiS Observational Study (n=1,939) and an interventional trial (n=519) at the same study sites with the same eligibility criteria^25^. The score was a summary measure of household wealth based on ownership of the following items: electricity, almirah, fan, table, chair, radio, television, fridge, pump, freezer, mobile, watch, computer, phone, animals, autobike, bicycle, rickshaw and a vehicle, where higher scores correspond to greater wealth. The resulting distribution was split into quintiles (Q1-Q5), leading to an uneven distribution of subjects in each quintile for our study cohort. For statistical analysis, the asset index was converted into a binary variable by grouping the lowest quintiles (Q1-Q2) and the highest quintiles (Q3-Q5). For maternal milk, infant age and maternal secretor status were the primary variables of interest. We also assessed metabolic changes across lactation stages, by grouping milk samples into colostrum (< 5 days), transitional (5-15 days), and mature milk (> 15 days).

### Untargeted metabolomics sample preparation

Metabolomics analysis was conducted on 380 infant fecal samples, 34 infant plasma samples, and 119 mother’s milk samples. Stool and plasma were extracted using 95% ethanol (EtOH) via the Matrix tube method as previously described^96,97^. Briefly, samples were extracted in 400 µL of 95% EtOH, shaked at 1,200 rpm for 2 min in a SpexMiniG plate shaker (SPEX SamplePrep part #1600, NJ, USA), and centrifuged for 5 min at 2,700 g. Supernatant (200 µL) was then collected and stored at -80°C for later acquisition. Prior to instrumental analysis, plates were redissolved in 200 µL 50:50 methanol (MeOH):H20. Milk samples were extracted in 75:25 MeOH:methyl tert-butyl ether (MTBE) as previously described^98^. Briefly, 400 µL of the MeOH/MTBE solution was added to 50 µL of milk, sonicated for 5 min, followed by a 15 min centrifugation (4°C) at 1800 rpm. Supernatant (300 µL) was immediately transferred to a new 96 well plate for instrumental analysis.

### Untargeted metabolomics data acquisition

Fecal and plasma samples were randomized and run on the same batch, while mother’s milk samples were run on a separate batch. Samples (5 μL) were injected into an UltiMate 3000 liquid chromatography system (Thermo Scientific) coupled to a QExactive Orbitrap (Thermo Scientific) mass spectrometer. Chromatography was conducted using a Kinetex C18 column (Phenomenex). A representative linear gradient of mobile phase A (water + 0.1% formic acid) and phase B (acetonitrile + 0.1% formic acid) consisted of 0-1 min isocratic at 5% B, 1-7 min to 98% B, 7-7.5 min isocratic at 98% B, 7.5-8 min to 5% B, and 8-10 min at 5% B with a flow rate of 0.5 mL/min. All solvents used were LC-MS grade. Data was acquired in data-dependent acquisition (DDA) mode using tandem mass spectrometry (MS/MS) in positive electrospray ionization (ESI+). Briefly, full scan MS spectra was acquired at 35,000 resolution with an automatic gain control (AGC) target of 5e5, maximum ion injection time of 100 ms, and a scan range of 100-1500 *m/z*. MS/MS spectra were collected using the same resolution and AGC target, and fragmented the top 5 most abundant ions per cycle with a 3.0 *m/z* isolation window, stepped normalized collision energies of 20, 30, and 40%, and a dynamic exclusion window of 10 s.

### Untargeted metabolomics data processing

Acquired .raw files were converted into .mzML format using ProteoWizard MSConvert^99^ and deposited in [GNPS/MassIVE](https://massive.ucsd.edu/ProteoSAFe/static/massive.jsp) under the accession code [MSV000096943](https://massive.ucsd.edu/ProteoSAFe/dataset.jsp?task=b5b8f8469ac74836a59fcc460b2ebed8). Feature detection and extraction were performed using MZmine 4.2^100^. Briefly, data were imported with MS1 and MS2 detector noise factor set to 3 and 2 respectively. The chromatogram builder parameters were set at 5 minimum consecutive scans, 1e5 minimum absolute height, and 10 ppm for *m/z* tolerance. Smoothing and local minimum feature resolver were applied, with the following parameters for the latter: chromatographic threshold 85%, minimum search range retention time 0.2 min, and minimum ratio of peak top/edge 2. Isotope filter and finder were then applied before feature alignment via join aligner (3 weight for m/z, 0.3 retention time tolerance). Peak finder was performed after removing features that were not detected in at least 3 samples. Further, metaCorrelate and ion identity molecular networking were run. List of MS/MS spectra and associated feature table was then exported and used to generate a feature based molecular network (FBMN)^101^ on GNPS2^102^. The FBMN job is publicly available at https://gnps2.org/status?task=d3c90a3743cf435291b635404821ab53. Briefly, fragment tolerances for both parent and fragment ions were set at 0.02 and networking and annotation parameters were both set to 5 minimum matching peaks and cosine similarity > 0.7. The network.graphml file from the FBMN was imported and manipulated in Cytoscape^103^ (version 3.10.3). MS/MS spectra of unknown features were also run in SIRIUS^61^ (version 6.1.1) and CANOPUS^62^ was used to predict features’ chemical pathway, superclass or class. Features of interest were searched using microbeMASST^55^ (https://masst.gnps2.org/microbemasst/) (minimum 4 matching peaks and cosine similarity > 0.7; output from 07/15/2025), a taxonomically informed mass spectrometry search tool within GNPS where MS/MS spectra can be linked to their respective microbial producers by querying against a curated reference database of >60,000 microbial monocultures.

### Untargeted metabolomics data analysis

The feature and annotation table were imported in R 4.4.2 (R Foundation for Statistical Computing, Vienna, Austria) for downstream data analysis. Sample’s total extracted peak areas were investigated to look into poor data acquisition due to sample collection, degradation, or instrument variation. Features with a retention time (RT) < 0.2 or > 8 min were excluded. Blank subtraction was performed by removing features detected in Blank samples meaning peak areas were not at least 5 times that observed in the samples. The package ‘homologueDiscoverer v 0.0.0.9000’ was used to remove polymers detected in the LC-MS/MS run^104^. Three fecal samples were excluded from the metabolomics analysis due to missing metadata, unusually high intensity in the ion total chromatogram, and clustering with the plasma samples. Features with near zero variance were removed using the package ‘caret v 6.0’. Features annotated as bile acid candidates using the propagated bile acid library^30^ underwent post-validation filtering based on the presence of diagnostic MS/MS fragment ions using massQL^105^ (https://massqlpostmn.gnps2.org/). Features were robust center log ratio (RCLR) transformed using ‘vegan v 2.6’ before applying dimensionality reduction techniques. Principal component analysis (PCA) and partial least square discriminant analysis (PLS-DA) models were built using ‘mixOmics v 6.22’. PERMANOVA was used to test centroid separation. The performances of the PLS-DA models were investigated via a 5-folds cross-validation. VIP scores > 1 were considered significant for group separation. The natural log ratios were generated by summing the peak areas of features enriched in one condition (numerator) relative to features enriched in the other condition (denominator). For compounds with multiple annotations, e.g. due to isomers producing similar MS/MS, peak areas were summed when plotting their trajectories over time (e.g. this was the case for some *N*-acyl lipids). The ‘lmerTest v 3.1’ package was used to generate the linear mixed effect models, where subject id was used as random effect and infant age as a fixed effect. For panels showing trajectories with age, smoothed trend lines using ‘LOESS’ (non-parametric local regression smoothing method) were added for visualization purposes, and were not derived from LME. The Wilcoxon rank-sum test was used to determine statistically significant group differences followed by Benjamini-Hochberg correction with p value < 0.05 considered statistically significant. To avoid repeated measures in comparisons across time windows (e.g. days 0-7, 14-28, and 30-45), only the sample with the highest value (peak area or log ratio) per subject within each window was retained. For the subanalysis investigating discriminant features between days 10-30 for delivery mode, one fecal sample per infant was selected, specifically the sample with the highest log ratio within this interval. Features detected in > 20% of the samples were included. Log₂ fold changes were calculated between groups and features with an FDR-adjusted p value < 0.10 were considered relevant. Upset plots were generated using the package ‘UpSetR v 1.4’^106^. Only human milk from Secretors (functional *FUT2* gene) was included in the correlation analysis between the targeted quantification of HMOs using HPLC after fluorescent derivatization and the untargeted LC-MS/MS data. Zero values, only present in the untargeted data, were also excluded from the analysis (46% of samples for 2’FL and 54% of samples for DFLac).

### Sample processing for metagenomic sequencing

All samples were transferred into 1 mL Matrix Tubes (ThermoFisher, Waltham, MA, USA) for gDNA extraction. Samples were extracted for metagenomic sequencing using reagents from the MagMAX Microbiome Ultra Nucleic Acid Isolation Kit (ThermoFisher Scientific, Waltham, MA, USA) following a previously described high-throughput protocol^107^, minimizing reaction volumes, eliminating the need for the sample-to-bead plate transfer step, and reducing potential cross-contamination from vortexing during bead beating. Extracted gDNA was quantified using the Quant-iT PicoGreen dsDNA Assay Kit (Invitrogen, Waltham, MA, USA) and normalized to 5 ng in 3.5 µL sterile water for library preparation, which was performed using a miniaturized adaptation of the KAPA HyperPlus Library Kit (Roche, Basel, Switzerland)^108^.

The library was quantified via PicoGreen Assay (Invitrogen), and all samples were equal volume pooled, polymerase chain reaction (PCR) cleaned (Qiagen), and size selected from 300 to 700 bp using a Pippin HT (Sage Sciences). QC was run on an Agilent 4000 Tapestation (Agilent, Santa Clara, CA, USA) to confirm expected library sizes after PCR cleanup and size selection. The equal volume pool was sequenced on an iSeq100 (Illumina, San Diego, CA, USA). Utilizing the sample concentration and read counts per sample obtained from the iSeq 100 run, a normalized pooling value was calculated for each sample to optimize pooling efficiency to obtain more even read counts per sample during NovaSeq sequencing^109^. After re-pooling the library with the iSeq normalized pool values, samples were PCR cleaned, size selected (300–700 bp), and QC was performed using an Agilent tapestation.

### Sample bioinformatic analysis

The PCR-cleaned, size-selected, iSeq-normalized pool was sequenced on a NovaSeq 6000 (Illumina, San Diego, CA, USA) at the Institute for Genomic Medicine at the University of California, San Diego with an S4 flow cell and 2 × 150 bp chemistry. Raw sequence reads (BCL files) were demultiplexed to per sample FASTQ and quality filtered^110^. Adapter trimming was performed by fastp^111^, and human reads were filtered out to ensure compliance with database regulations. This was performed via the conservative host filtration pipeline described previously^112^. Briefly, reads were first filtered using minimap2^113^ via alignment to two reference genomes: GRCh38.p14 plus PhiX, and CHM13v2.0. Additionally, a pangenome index was created using the 94 currently available pangenomes from the Human Pangenome Reference Consortium^114^ using Movi^115^. Unmapped reads following alignment were queried against this index, and reads with high-propensity matches to one or more human pangenome references were discarded. The resulting FASTQ files were uploaded into Qiita^116^ (study ID #14834) and processed using the Woltka pipeline^117^ (version 0.1.7) using SHOGUN^118^ parameters. In brief, direct genome alignments were made against the “Web of Life” database (release 2)^119^, which contains 15,953 microbial genomes of bacteria and archaea. Paired-end sequence alignment is performed using the bowtie2 (v2.5.4) aligner^120^ (using --very-sensitive -k 16 --np 1 --mp "1,1" -- rdg "0,1" --rfg "0,1" --score-min "L,0,-0.05" --no-head --no-unal --no-exact-upfront --no-1mm- upfront) and by mapping sequencing data to microbial reference genomes^118^. Microbial genome IDs are considered operational genomic units (OGUs)^117^. The OGU frequencies were then summed after the entire alignment was processed and rounded to the nearest even integer, thereby making the sum of OGU frequencies per sample nearly equal (considering rounding) to the number of aligned sequences in the data set. The resultant count matrix is saved and formatted as a BIOM format table^121^, which has microbial genome IDs (rows) by samples (columns) that contains read counts as values.

The BIOM file was converted to a QIIME2^122^ (v2024.5) artifact. We applied a novel coverage dispersion filter called micov which eliminated OGU with less than 25% reference genome coverage (https://github.com/biocore/micov) to reduce risk of false positives from short read multimapping. A 1% prevalence filter was also applied. Contamination in the resultant table was removed using SCRuB (Source-tracking for Contamination Removal in microBiomes)^123^, a probabilistic decontamination method that incorporates samples and controls to remove contamination. Due to the large sample size of this study SCRuB was run separately for each different sample plate to account for unique well-to-well contamination. The resulting table contained 1,568,600,606 reads for 1,609 samples, with an average of 901,495 reads per sample. Standard alpha diversity metrics (Faith’s PD^124^, Shannon index^125^), as well as standard beta diversity metric, unweighted UniFrac^126^, and phylogenetic robust center-log-ratio (rclr) metric, phylo-RPCA^36^, were calculated. For unweighted UniFrac, dimensionality was reduced using PCoA and visualized using EMPeror^127^. For phylo-RPCA, dimensionality was reduced using PCA and visualized using EMPeror. For unweighted UniFrac, the feature table was rarified according to a read count value that would retain 98.5% of all samples from all body sites (including 90% of vaginal samples), 644 reads. Alpha diversity rarefaction depth for low biomass samples (vaginal, human milk) is 644, high biomass samples is 100,000 (fecal, oral, skin). For rclr-based diversity metrics (phylo-RPCA), an unrarefied table (sample minimum read count for inclusion was 644 reads for cross body site comparison in Extended Data Fig. 4. When binned timepoints are used, samples are grouped into Day 0, 7, 14, 30, 60, 90, 180 to which they are closest. TEMPoral TEnsor Decomposition (TEMPTED)^128^ is a beta diversity dimensionality reduction method that accounts for repeated sampling and uses time as a continuous variable, plus it can handle varying temporal sampling present in the dataset.

Beta diversity differences were determined by PERMANOVA. Feature tables were collapsed to the genera level and then differential abundance was performed using analysis of compositions of microbiomes with bias correction (ANCOM-BC2)^129^. ANCOM-BC2 utilized the binary age category (greater than 1 month vs. less than 1 month of age) in the differential abundance modeling formula. The differentially abundant genera were used to create a natural log ratio of the sum of all read counts from genera enriched after one month (numerator) to all genera depleted (denominator). Qurro was used to visualize ranked features^130^. ConvexHull (CH) volume and area are measures that capture the dispersion or spread of a set of points in a multidimensional space (scipy ConvexHull v0.12.0). CH volume has been used in the past to assess the stability of microbial communities over time^131^. Here, a convex hull volume or area is calculated over all samples of a given body site at a given time. We also employed several linear mixed-effects models (LME) using the statsmodels package in Python to assess whether dependent variables of interest (faith pd values, convex hull volumes, etc.) varies with fixed effects. Generally, infant age at time of sample (days postpartum) was used as a fixed effect and individual host subject identification was used to account for repeated measures. For example, most of the alpha diversity metrics’ LME formulas were “faith_pd ∼ host_age_infant + (1 | host_subject_id)”. For **Fig 1h**, the formula was “ln(*bsh* reads) ∼ ln(AA/Tau conjugates) + (1 | host_subject_id)”. For **Extended Data Fig. 6f**, the formula was “[faith_pd or log_ratio] ∼ host_age_infant * hmo_secretor + (1 | host_subject_id)”, the star indicates an interaction term and is the one reported for significance. For panels showing trajectories with age, lineplots using binned timepoints for smoothing were added for visualization purposes and were not derived from LME. When binned timepoints were used in boxplots, differences were assessed using two-sided Mann-Whitney-Wilcoxon, followed by Holm-Bonferroni correction to adjust for multiple comparisons (2MWW-HB).

Fast expectation-maximization microbial source tracking (FEAST) was used to estimate the contribution of potential source environments to a given sample^38^. Infant fecal samples were used as sinks in this analysis. The early timepoint used was from the birth/0 day sample, or if unavailable, the first sample collected for each infant. The late timepoint used the 6 month/180 day sample, or if unavailable, the final sample collected for each infant. For each timepoint dataset, we examined the estimated contribution proportion of matched or unmatched maternal biospecimens as potential sources. Using different_sources_flag = 0, all mother vaginal, fecal, and milk samples across all timepoints are given as potential sources. Using different_sources_flag = 1, all paired-mother vaginal, fecal, and milk samples across all timepoints are given as potential sources.

### Bile salt hydrolase/transferase gene search

Gene searches were specifically performed for bile acid modification enzymes *(hsdh, bai, bs*h). For each gene of interest, faa files of all matches were acquired from Uniprot^132^, pooled, and then amino acid sequences with at least 60% similar identity were clustered via USEARCH^133^ into a unique database. Then, metagenomic reads from each sample were aligned and matched to the genes of interest against the respective USEARCH clustered databases using Diamond blastx^134^. Output data frames were further filtered for matches with >40 bp of alignment length, > 50% query alignment ratio, and e-value< 0.00001 before plotting.

### Multi-omics analysis

Joint-RPCA is an unsupervised method used to agnostically determine co-occurrences between the microbiome and metabolome data^36^. Since the method is not optimized for longitudinal sampling, samples around day 20 (day 10-30) were chosen for analysis. In short, the raw features tables for each method after previously described filtration methods were robust center log ratio (RCLR) transformed. RCLR of the raw observed values were only computed on the non-zero entries and then averaged and optimized. This helps to account for the sparsity and compositionality. The shared samples of all input matrices were used to estimate a shared matrix. The estimated shared matrix and the matrix of shared eigenvalues across all input matrices were recalculated at each iteration to ensure consistency. Minimization was performed across iterations by gradient descent. Cross-validation of the reconstruction was performed in order to prevent overfitting of the joint factorization. Co-occurrences of features across the two input matrices were calculated from the final estimated matrices.

### HMO analysis

HMO profiles were obtained from 130 samples from 55 mothers at enrollment (0/1), day 60 and 2 other timepoints up to day 60 of the infant’s age. HMOs were analyzed using HPLC after fluorescent derivatization allowing for quantification of the 19 most abundant HMOs as previously described^135,136^. HMO concentrations are reported in nmol/mL, and the sum of all HMOs in a sample was calculated as the sum of all HMOs detected in each sample. HMO-bound sialic acid and HMO-bound fucose were calculated as the sum of sialic acid and fucose moieties included in each HMO. Overall, 19 HMOs were identified and quantified: 2′-fucosyllactose (2′FL), 3-fucosyllactose (3FL), 3′-sialyllactose (3′SL), 6′-sialyllactose (6′SL), difucosyllactose (DFLac), difucosyllacto-*N*-hexaose (DFLNH), difucosyllacto-*N*-tetrose (DFLNT), disialyllacto-*N*-hexaose (DSLNH), disialyllacto-*N*-tetraose (DSLNT), fucodisialyllacto-*N*-hexaose (FDSLNH), fucosyllacto- *N*-hexaose (FLNH), lacto-*N*-fucopentaose (LNFP) I, LNFP II, LNFP III, lacto-*N*-hexaose (LNH), lacto-*N*-neotetraose (LNnT), lacto-*N*-tetrose (LNT), sialyl-lacto-*N*-tetraose b (LSTb), and sialyl- lacto-*N*-tetraose c (LSTc).

## Data availability

Untargeted LC-MS/MS data generated for this study are publicly available at GNPS/MassIVE (https://massive.ucsd.edu/) under the accession code [MSV000096943](https://massive.ucsd.edu/ProteoSAFe/dataset.jsp?task=b5b8f8469ac74836a59fcc460b2ebed8). The associated FBMN job is publicly available at GNPS2: https://gnps2.org/status?task=d3c90a3743cf435291b635404821ab53. Metagenomics sequencing data will be available through EBI/ENA via the following accession codes: PRJEB83236 ERP166886 (on publication). Additional information and processing pipelines will be available in Qiita Study ID #14834 (https://qiita.ucsd.edu/study/description/14834).

## Code Availability

Metagenomics analyses located at https://github.com/knightlab-analyses/gates_sepsis_bangladesh.

Metabolomics analyses located at https://github.com/kinekvitne/manuscript_gates_sepsis_obs

## Acknowledgements

The authors would like to acknowledge the work of Gail Ackermann, who assisted with project management at UC-San Diego.

## Author contributions

K.E.K. conducted untargeted metabolomics analysis, multi-omics analysis, generated figures, and wrote the manuscript.

C.A. conducted metagenomics analysis, multi-omics analysis, generated figures, and wrote the manuscript.

J.C.O was involved in metadata preparation, conducted metagenomics analyses, and wrote sections of the manuscript.

D.P. conducted metagenomics analysis and edited the manuscript.

S.Z., V.C.L., I.M., and A.P. performed untargeted metabolomics analysis.

S.T. processed untargeted metabolomics samples.

K.S., L.P.,and L.B. conducted HMO analysis.

A.G. and D.M. processed metagenomics samples.

L.A.B. provided feedback on analysis, wrote portions of manuscript and edited manuscript.

S.M.T. provided feedback and secured funding.

L.G.P and D.E.R. designed the SEPSiS project from which data were used for the present study; provided feedback on the manuscript.

L.B., P.C.D. and R.K. designed the study, provided feedback, and secured funding. All authors reviewed and approved the manuscript.

## SUPPLEMENTARY TABLE TITLES AND LEGENDS

**Supplementary Table 1.** Output from the PLS-DA models showing variable importance in projection (VIP) scores across variables for maternal milk and infant feces: lactation stage (maternal milk), maternal secretor status (maternal milk and infant feces), delivery mode (infant feces), and household drinking water treatment (maternal milk and infant feces).

**Supplementary Table 2.** Output from multivariate PERMANOVA.

**Supplementary Table 3.** Overlapping discriminant features for water treatment identified by PLS-DA.

