## Supplementary Figures for "Environmental and Maternal Imprints on Infant Gut Metabolic Programming"

[1] Skaggs School of Pharmacy and Pharmaceutical Sciences, University of California San Diego, La Jolla, CA, USA
[2] Department of Pediatrics, University of California San Diego, La Jolla, CA, USA
[3] Centre for Global Child Health, The Hospital for Sick Children, Toronto, Ontario, Canada
[4] Department of Nutritional Sciences, Faculty of Medicine, University of Toronto, Toronto, Ontario, Canada
[5] Bioinformatics and Systems Biology Program, University of California San Diego, La Jolla, CA, USA
[6] Collaborative Mass Spectrometry Innovation Center, University of California San Diego, La Jolla, CA, USA
[7] Medical Scientist Training Program, University of California, San Diego, La Jolla, CA, USA
[8] Larsson-Rosenquist Foundation Mother-Milk-Infant Center of Research Excellence (MOMI CORE), University of California San Diego, La Jolla, CA, USA
[9] Human Milk Institute (HMI), University of California San Diego, La Jolla, CA, USA
[10] Nutrition Research Division, International Centre for Diarrhoeal Disease Research, Bangladesh, Dhaka, Bangladesh
[11] Shield Pharmaceuticals Corp., Ronkonkoma, NY, USA
[12] Child Health Evaluative Sciences, The Hospital for Sick Children, Toronto, Ontario, Canada
[13] Dalla Lana School of Public Health, University of Toronto, Toronto, Ontario, Canada
[14] Department of Global Health, Boston University School of Public Health, Boston, MA, USA
[15] Department of Medicine, Boston University Chobanian & Avedisian School of Medicine, Boston, MA, USA
[16] Child Health Research Foundation, Dhaka, Bangladesh
[17] Department of Paediatrics, Faculty of Medicine, University of Toronto, Toronto, Ontario, Canada
[18] Department of Pediatrics, Mt. Sinai Hospital, Toronto, Ontario, Canada
[19] Mohammadpur Fertility Services and Training Centre, Dhaka, Bangladesh
[20] Department of Obstetrics Gynecology and Reproductive Sciences, University of California San Diego, La Jolla, CA, USA
[21] Center for Microbiome Innovation, University of California San Diego, La Jolla, CA, USA
[22] Department of Computer Science and Engineering, University of California San Diego, La Jolla, CA, USA
[23] Shu Chien-Gene Lay Department of Bioengineering, University of California San Diego, La Jolla, CA, USA
[24] Halıcıoğlu Data Science Institute, University of California San Diego, La Jolla, CA, USA
* Contributed equally

### EXTENDED DATA FIGURES

a

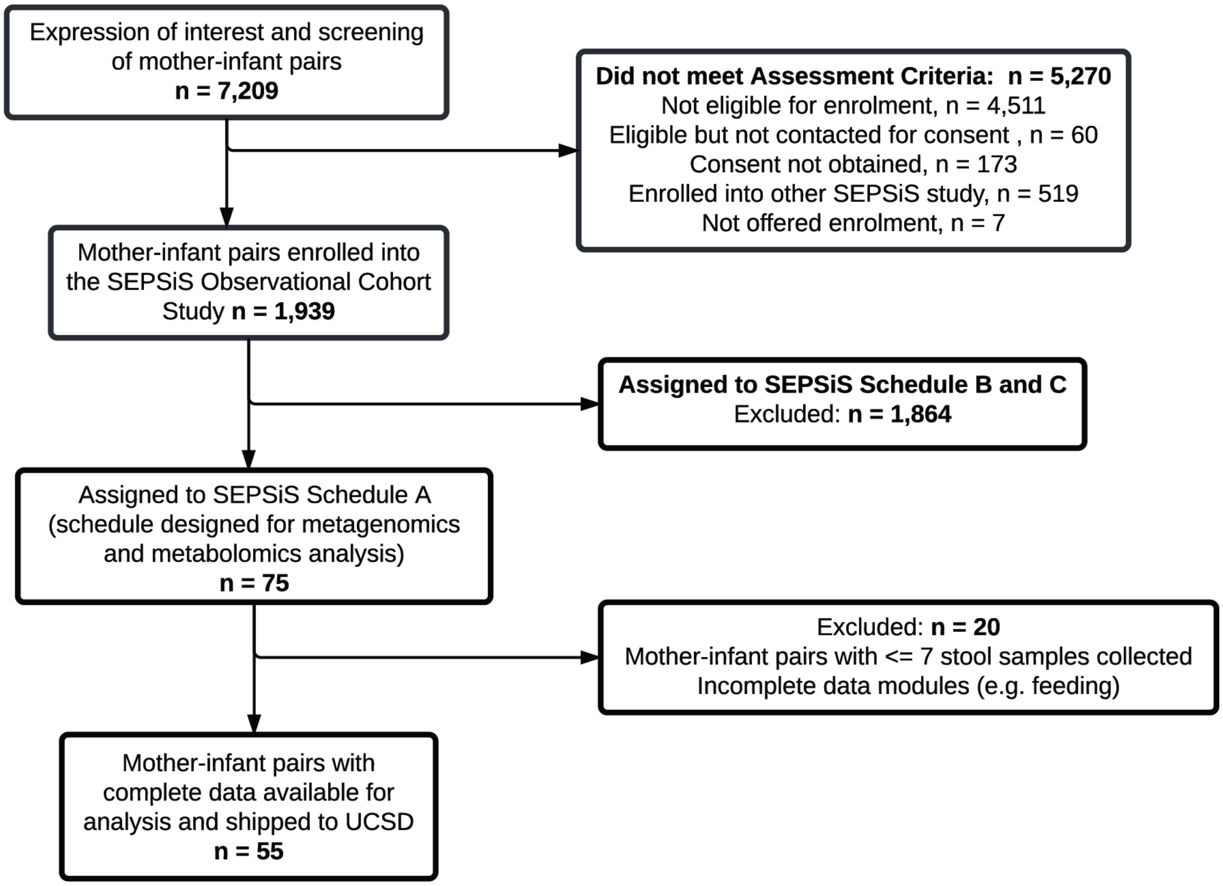


**Extended Data Fig. 1: Study flow diagram. a,** Detailed flowchart for participant and sample selection.

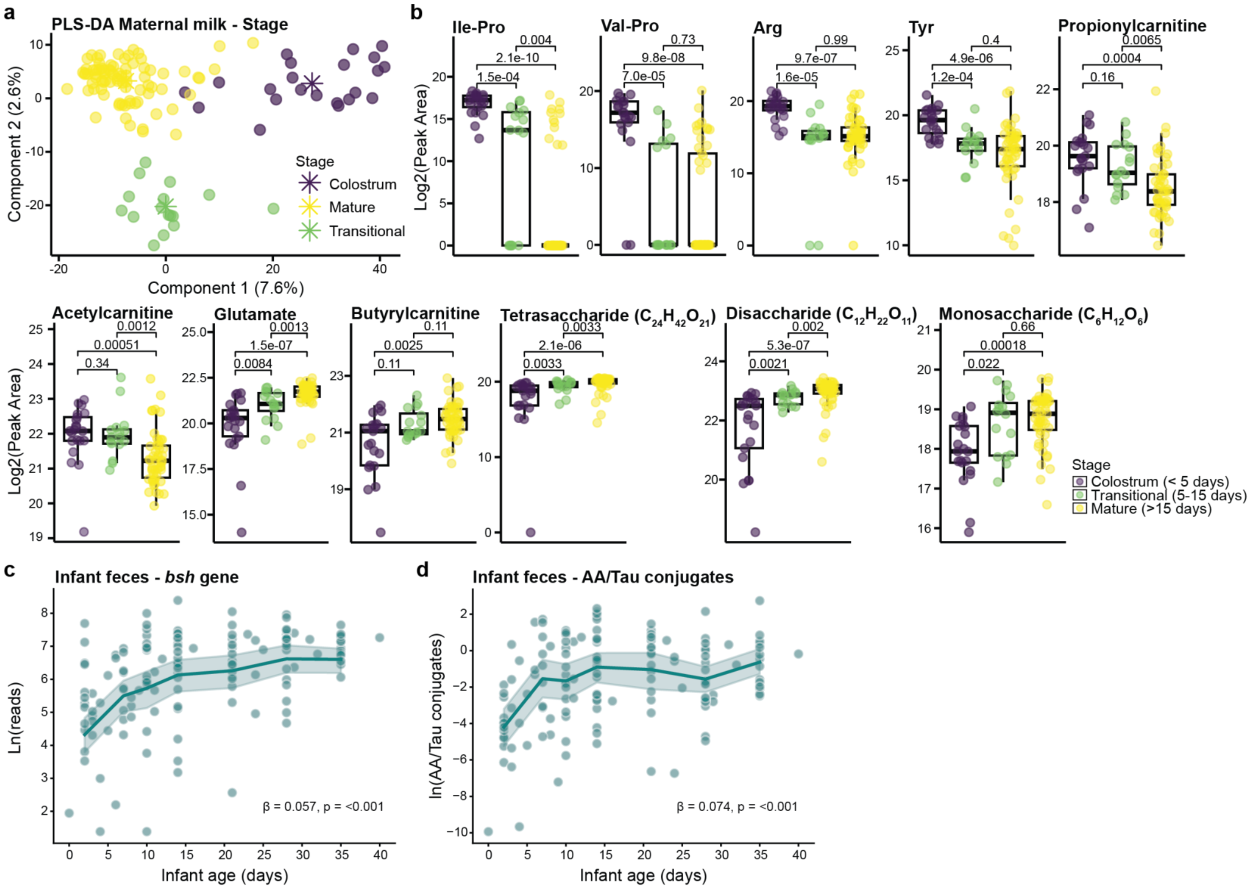


**Extended Data Fig. 2: The maternal milk metabolome changes across lactation changes. a,** Supervised PLS-DA of maternal milk samples by lactation stage (classification error rate (CER) = 0.18). **b,** Boxplot showing the natural log peak areas of key discriminant features across lactation stages in human milk. Statistical significance was determined by the Wilcoxon test with Benjamini-Hochberg correction for multiple comparisons. Repeated measures were avoided by retaining only the highest value per subject within each lactation stage. All boxplots show the first (lower), median, and third (upper) quartiles, with whiskers 1.5 times the interquartile range. **c,** Trajectory of the natural log ratio of bile salt hydrolase/transferase (*bsh*) gene reads in infant feces with increasing infant age; lineplot used binned timepoints. **d,** Trajectory for the natural log ratio of amino acid over taurine conjugated bile acids in early life; lineplot used binned timepoints. β and p values in **c** and **d** were derived from linear mixed effects models (LME) with subject as a random effect.

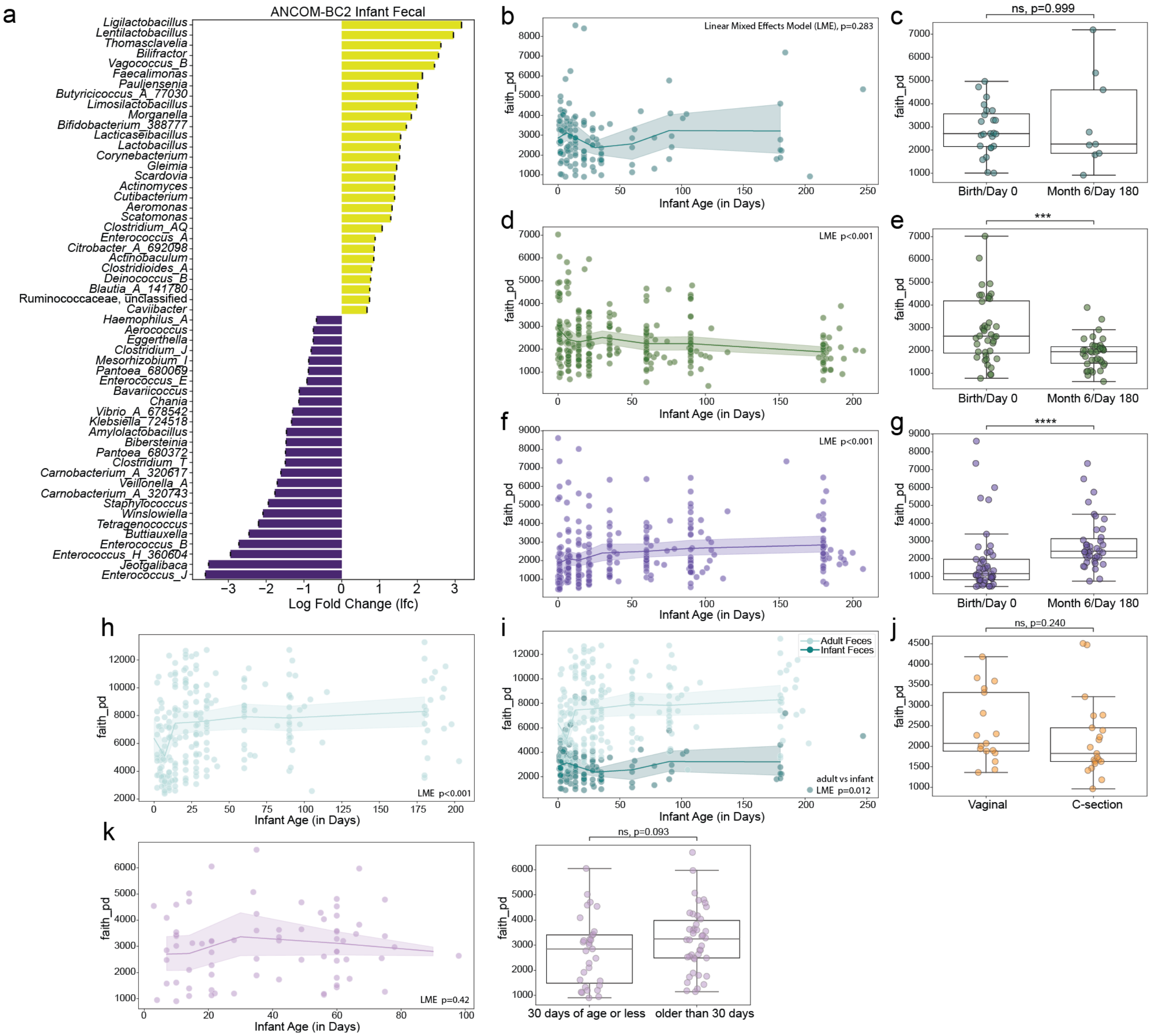


**Extended Data Fig. 3: Cross Body Site Metagenomics Alpha Diversity. a,** Differentially abundant infant fecal genera before and after 30 days of age as identified by ANCOM-BC2 ($p < 0.05$, LFC > 0.5). This is the full list of genera used in the creation of the log ratio**.** Faith's Phylogenetic Alpha Diversity: each body site is rarefied to a unique depth that retains 90% of samples (see Methods for number). Lineplots use binned timepoints for smoothing. **b,** Infant feces longitudinal scatterplot with lineplot. **c,** Infant feces boxplot with binned timepoints, Birth/Day 0 compared to Month 6/Day 180. **d,** Infant skin longitudinal scatterplot with lineplot. **e,** Infant skin boxplot with Day 0 compared to Day 180. **f,** Infant oral (tongue) longitudinal scatterplot with lineplot. **g,** Infant oral boxplot Day 0 compared to Day 180. **h,** Adult feces longitudinal scatterplot with lineplot. **i,** Adult and infant feces longitudinal scatterplot with lineplot. **j,** Adult vaginal (only sampled at Day 0/birth of infant) boxplot comparing Vaginal and C-section. Significance determined by LME with interactions, formula = "faith_pd ~ host_age_infant * host_life_stage + (1 | host_subject_id)". The coefficient and p-value is for the interaction term. **k,** [Left] Human milk longitudinal scatterplot with lineplot. [Right] Human milk boxplot comparing early (<30 days) to later (>30days) timepoints. Longitudinal significance determined by linear mixed effects model (LME). Boxplot significance determined by two-sided Mann-Whitney-Wilcoxon test. Notation: ns $p > 0.05$, * $p < 0.05$, ** $p < 0.01$, *** $p < 0.001$, **** $p < 0.0001$.

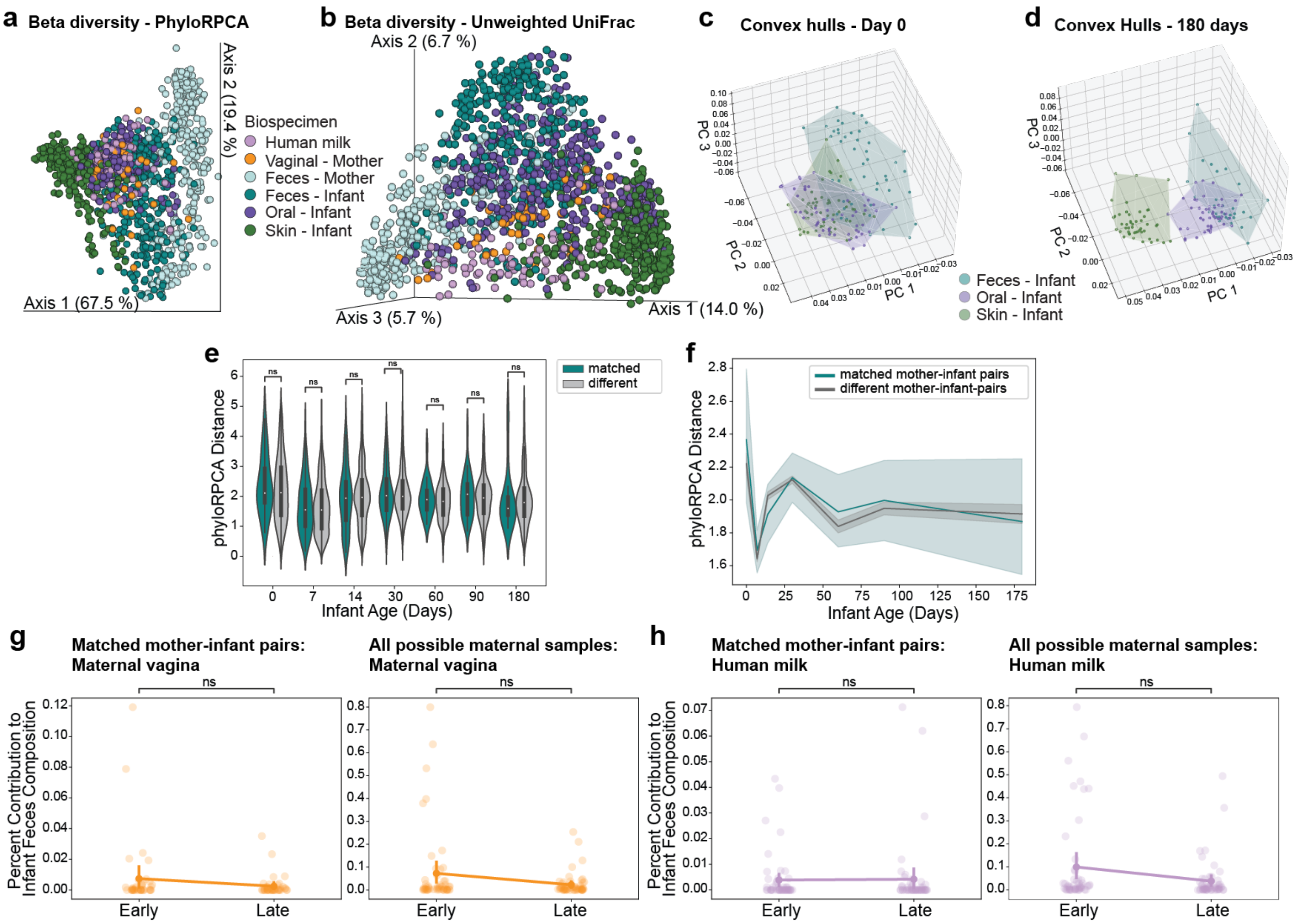


**Extended Data Fig. 4: Cross Body Site Metagenomics Beta Diversity and additional FEAST Source Tracking.** Beta Diversity: **a**, phyloRPCA PCA plot (samples less than 644 reads excluded). **b,** Unweighted UniFrac PCoA (rarefied 644). phyloRPCA ConvexHulls: **c,** Infant samples (feces, skin, oral/tongue) at Day 0/Birth. **d,** Infant samples Day 180/Month 6. **e,** Violinplots of phyloRPCA mother-infant fecal sample beta diversity distances that compare within matched mother-infant pairs and distances between different (unmatched) mothers and infants. The internal boxplot shows median in white and interquartile range. **f,** phyloRPCA lineplot of within and between mother-infant pair distances over time. Darker line indicates mean, shaded region indicates 95% confidence interval. FEAST: **g,** Stripplot/pointplot of the FEAST estimated percentage contribution of maternal vagina samples to matched (left) or all available (right) infant fecal samples. **h,** Stripplot/pointplot of the FEAST estimated percentage contribution of human milk samples to matched (left) or all available (right) infant fecal samples. Significance determined by two-sided Mann-Whitney-Wilcoxon test with Holm-Bonferroni correction. Notation: ns is not significant, $p > 0.05$.

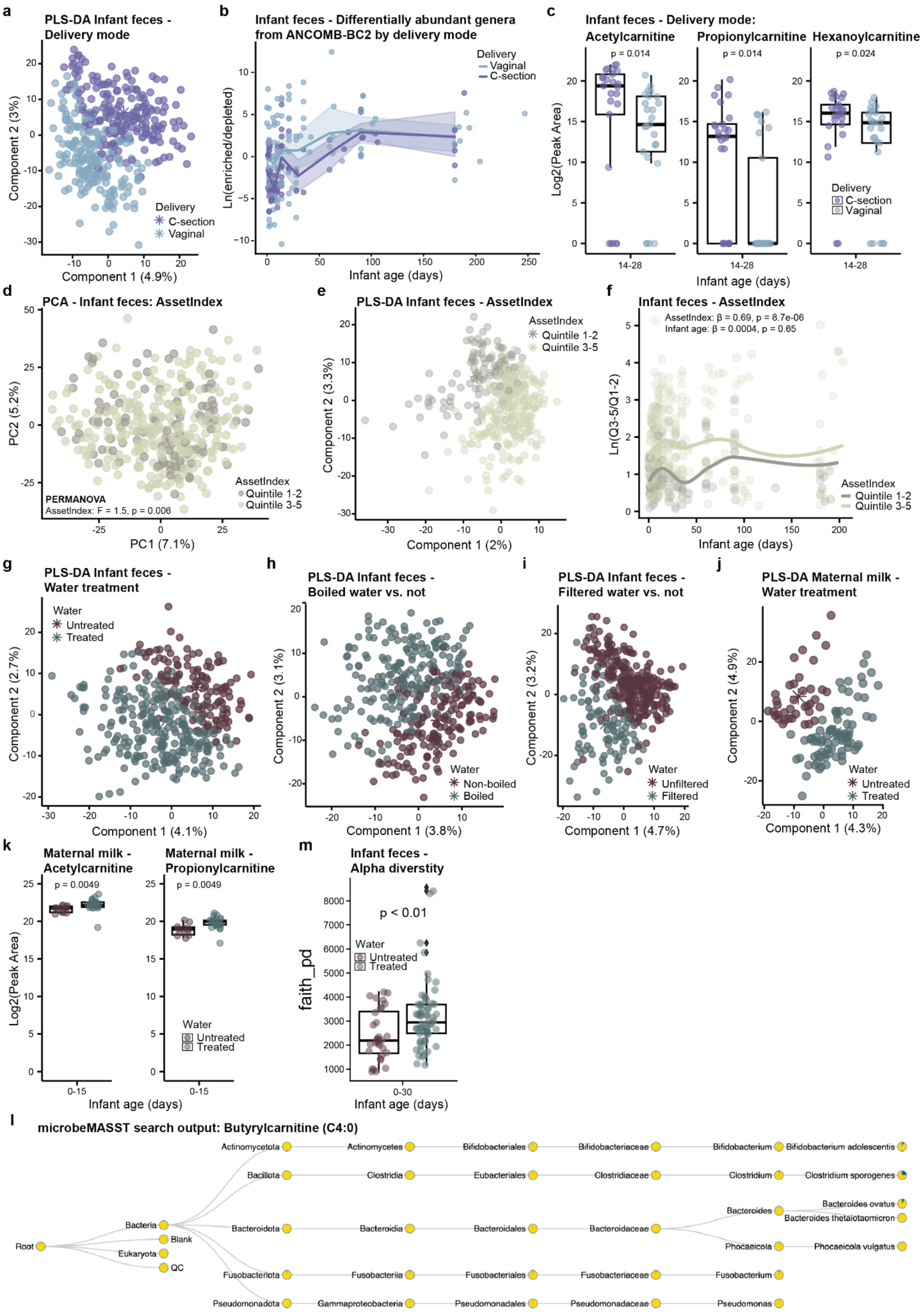
a
PLS-DA Infant feces - Delivery mode
Component 2 (3%)
Component 1 (4.9%)
Delivery
C-section
Vaginal
b
Infant feces - Differentially abundant genera from ANCOMB-BC2 by delivery mode
Ln(enriched/depleted)
Infant age (days)
Delivery
Vaginal
C-section
c
Infant feces - Delivery mode:
Acetylcarnitine
Propionylcarnitine
Hexanoylcarnitine
p = 0.014
p = 0.014
p = 0.024
Log2(Peak Area)
14-28
Infant age (days)
Delivery
C-section
Vaginal
d
PCA - Infant feces: AssetIndex
PC2 (5.2%)
PC1 (7.1%)
AssetIndex
Quintile 1-2
Quintile 3-5
PERMANOVA
AssetIndex: F = 1.5, p = 0.006
e
PLS-DA Infant feces - AssetIndex
Component 2 (3.3%)
Component 1 (2%)
AssetIndex
Quintile 1-2
Quintile 3-5
f
Infant feces - AssetIndex
AssetIndex: β = 0.69, p = 8.7e-06
Infant age: β = 0.0004, p = 0.65
Ln(Q3-5/Q1-2)
Infant age (days)
AssetIndex
Quintile 1-2
Quintile 3-5
g
PLS-DA Infant feces - Water treatment
Water
Untreated
Treated
Component 2 (2.7%)
Component 1 (4.1%)
h
PLS-DA Infant feces - Boiled water vs. not
Component 2 (3.1%)
Component 1 (3.8%)
Water
Non-boiled
Boiled
i
PLS-DA Infant feces - Filtered water vs. not
Component 2 (3.2%)
Component 1 (4.7%)
Water
Unfiltered
Filtered
j
PLS-DA Maternal milk - Water treatment
Component 2 (4.9%)
Component 1 (4.3%)
Water
Untreated
Treated
k
Maternal milk - Acetylcarnitine
Maternal milk - Propionylcarnitine
p = 0.0049
p = 0.0049
Log2(Peak Area)
0-15
Infant age (days)
Water
Untreated
Treated
m
Infant feces - Alpha diverstity
p < 0.01
faith_pd
Water
Untreated
Treated
0-30
Infant age (days)
l
microbeMASST search output: Butyrylcarnitine (C4:0)
Root
Bacteria
Blank
Eukaryota
QC
Actinomycetota
Actinomycetes
Bifidobacteriales
Bifidobacteriaceae
Bifidobacterium
Bifidobacterium adolescentis
Bacillota
Clostridia
Eubacteriales
Clostridiaceae
Clostridium
Clostridium sporogenes
Bacteroidota
Bacteroidia
Bacteroidales
Bacteroidaceae
Bacteroides
Bacteroides ovatus
Bacteroides thetaiotaomicron
Phocaeicola
Phocaeicola vulgatus
Fusobacteriota
Fusobacteriia
Fusobacteriales
Fusobacteriaceae
Fusobacterium
Pseudomonadota
Gammaproteobacteria
Pseudomonadales
Pseudomonadaceae
Pseudomonas

**Extended Data Fig. 5: PLS-DA models and selected differential features in infant feces by delivery mode and water treatment. a,** Supervised PLS-DA of infant fecal samples by delivery mode (CER = 0.40). **b**, Natural-log ratio of enriched to depleted infant fecal genera (defined in Fig. 2b) by delivery mode. Overall, trajectories did not differ between C-section and vaginal delivery. Significance determined by LME with interactions, formula = "log_ratio ~ host_age_infant * mode_delivery + (1 | host_subject_id)". The p-value is for the interaction term (LME: p = 0.118), but there was a transient difference at Day 7 (2MWW-HB, p = 0.006). **c,** Boxplot showing differences in the natural log peak areas of short-chain acylcarnitines by delivery mode between Days 14-28. Statistical significance was determined by the Wilcoxon rank-sum test with BH adjusted p values. **d,** PCA of infant fecal samples showed separation by asset index, a proxy for socioeconomic status (PERMANOVA: p = 0.006). Households in quintile 1-2 were compared against quintile 3-5 (greater wealth) **e,** Supervised PLS-DA of infant fecal samples by asset index (CER = 0.28). **f,** Natural log ratios of the summed peak areas of the top 100 discriminating features for asset index. Statistical significant difference between groups was determined using LME with subject as random effect. Supervised PLS-DA of infant fecal samples by **g,** water treatment (CER = 0.35), **h,** boiled water vs. non-boiled water (CER = 0.44), and **i,** filtered water vs. unfiltered water (CER = 0.32). **j,** Supervised PLS-DA of human milk samples by water treatment (CER = 0.32). **k,** Boxplot showing differences in the natural log peak areas of selected acylcarnitines enriched in the untreated water group between days 0-15. In case of multiple samples per subject, only the highest value (peak area) was retained. Statistical significance was determined by the Wilcoxon rank-sum test with BH adjusted p values. **l,** microbeMASST search output for butyrylcarnitine (C4:0). The pie charts show the proportion of spectral matches; blue=match, yellow=non-match. **m,** Boxplot showing differences in Faith's phylogenetic alpha diversity between the treated versus untreated water group between days 0-30 (2MWW-HB, p = 0.0072). All boxplots show the first (lower), median, and third (upper) quartiles, with whiskers 1.5 times the interquartile range.

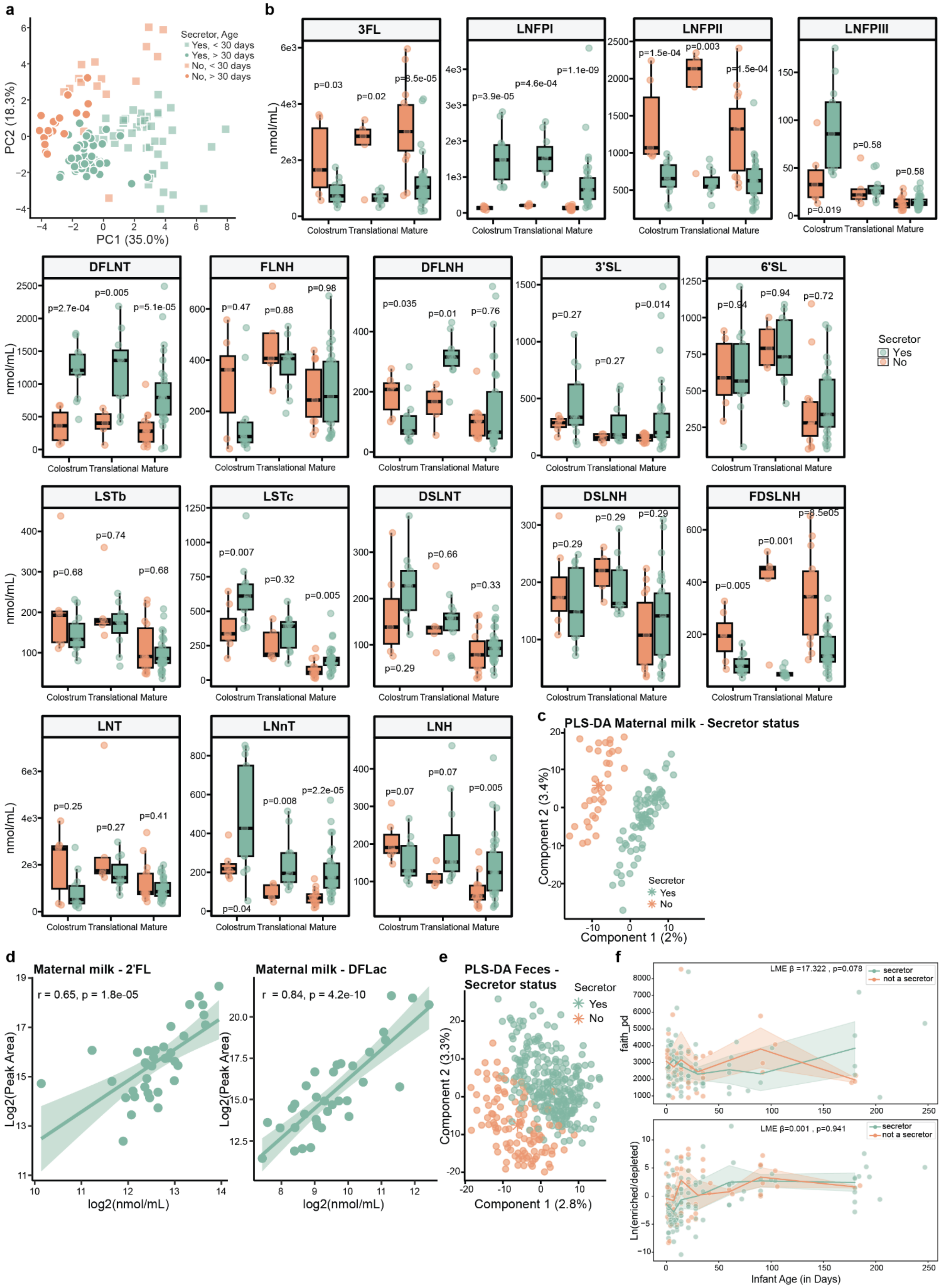
a
Secretor, Age
Yes, < 30 days
Yes, > 30 days
No, < 30 days
No, > 30 days
PC2 (18.3%)
PC1 (35.0%)
b
3FL
LNFPI
LNFPII
LNFPIII
DFLNT
FLNH
DFLNH
3'SL
6'SL
LSTb
LSTc
DSLNT
DSLNH
FDSLNH
LNT
LNnT
LNH
nmol/mL)
Colostrum Translational Mature
Secretor
Yes
No
c
PLS-DA Maternal milk - Secretor status
Component 2 (3.4%)
Component 1 (2%)
Secretor
Yes
No
d
Maternal milk - 2'FL
r = 0.65, p = 1.8e-05
Log2(Peak Area)
log2(nmol/mL)
Maternal milk - DFLac
r = 0.84, p = 4.2e-10
Log2(Peak Area)
log2(nmol/mL)
e
PLS-DA Feces - Secretor status
Secretor
Yes
No
Component 2 (3.3%)
Component 1 (2.8%)
f
LME β =17.322 , p=0.078
secretor
not a secretor
faith_pd
LME β=0.001 , p=0.941
secretor
not a secretor
Ln(enriched/depleted)
Infant Age (in Days)

**Extended Data Fig. 6. Human milk oligosaccharide (HMO) profiles differ based on secretor status.** Maternal milk samples (n=130) from 55 mothers were analyzed using high-performance liquid chromatography after fluorescent derivatization to quantify 19 various HMOs. Absolute concentration values are in nmol/mL. **a,** PCA of HMO profiles grouped by maternal secretor status and before/after 30 days postpartum (infant age). Mothers with unknown/undetermined secretor status were excluded. Significance was determined by PERMANOVA: by secretor status - Pseudo-F: 169.15, p-value: 0.0001; infant age group (before or after 30 days), Pseudo-F: 11.45, p-value: 0.0002; by secretor status and age group - Pseudo-F: 78.67, p-value: 0.0001. **b**, Boxplot showing quantified levels of fucosylated, sialylated, and neutral HMOs in human milk across lactation stages, grouped by secretor status (Wilcoxon rank-sum test with BH correction). In cases with multiple samples per subject within the stage, only the sample with the highest 2′FL concentration was retained. **c,** Supervised PLS-DA of maternal milk samples by secretor status (CER = 0.36). **d,** Scatterplot showing the correlation between targeted quantification and untargeted LC-MS/MS data for 2′-fucosyllactose (2′FL) and difucosyllactose (DFLac) (Pearson correlation). Only human milk from Secretors (functional *FUT2* gene) were included, and zero values were excluded from the analysis. **e,** Supervised PLS-DA of infant fecal samples by secretor status (CER = 0.34). **f,** [top] Longitudinal scatterplot with lineplot of Faith's PD alpha diversity metric for infant feces by secretor status. [bottom] Longitudinal scatterplot with lineplot of the previously created log ratio from ANCOM-BC2 results (**Fig. 3b,c; Extended Data Fig. 3a)** for infant feces by secretor status. Lineplots use binned timepoints. Significance determined by LME with interactions, the coefficient and p-value is for the interaction term.

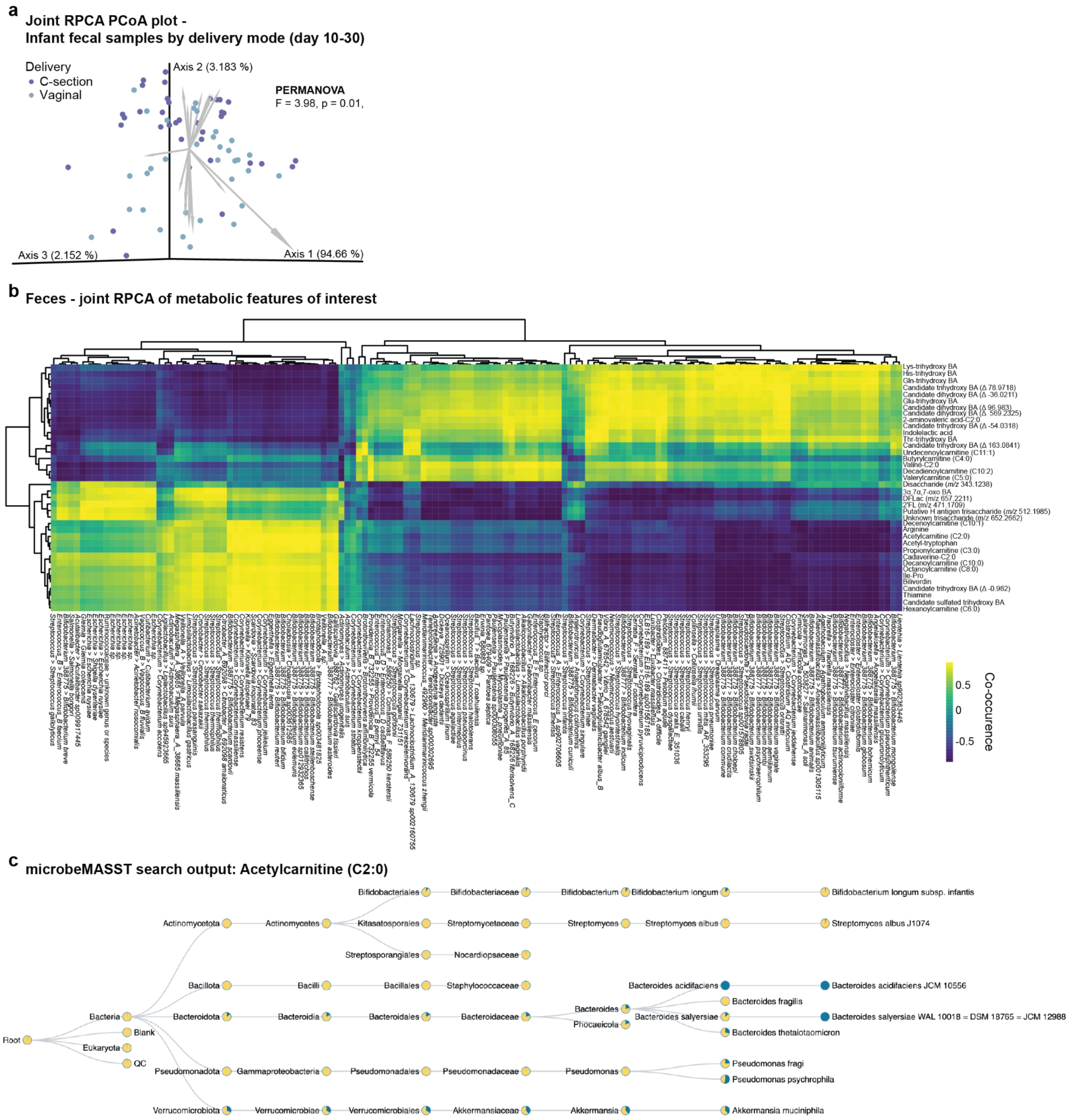


**Extended Data Fig. 7. Co-occurring metabolites and microbial features from a multi-omics analysis. a,** joint-RPCA PC1 vs PC2 vs PC3 are displayed in 3D using EMPeror. **b,** Heatmap of correlating metabolites and microbial species from infant fecal samples collected at day ~20 (between day 10-20) identified by joint-RPCA. Metabolic features shown were selected based on previous analyses. **c,** microbeMASST search output for acetylcarnitine (C2:0), with pie charts showing the proportion of spectral matches; blue=match, yellow=non-match.
