## Supplementary Table S2 for "Environmental and Maternal Imprints on Infant Gut Metabolic Programming"

**Supplementary Table 2. Output from multivariate PERMANOVA.** A permutational multivariate analysis of variance (PERMANOVA) was performed to investigate contribution of covariates to the variation in infant fecal metabolome. In addition to infant age, which was the primary variable of interest, the model also included other variables with known relevance to early life microbiome and metabolome.

| Variable | F value | R2 | P value |
|---|---|---|---|
| **infant_age_days** | **4.5** | **0.012** | **0.001** |
| **mode_delivery** | **1.9** | **0.005** | **0.001** |
| host_subject_id (accounted for in LME models) | 1.6 | 0.004 | 0.002 |
| **hmo_Secretor** [+] | **1.5** | **0.004** | **0.006** |
| **AssetIndex2*** | **1.5** | **0.004** | **0.006** |
| **water_treatment**[§] | **1.4** | **0.004** | **0.02** |
| gestational_age_birth | 1.4 | 0.004 | 0.014 |
| sex | 1.4 | 0.004 | 0.009 |
| birthweight | 1.2 | 0.003 | 0.091 |
| fp_long_bin[‖] | 1.2 | 0.003 | 0.102 |
| maternal_antibiotics[‡] | 1.1 | 0.003 | 0.287 |
| im_ever_abxs_adhoc[#] | 1.0 | 0.003 | 0.363 |

[+]*Secretors/non-secretors were defined by the presence or near-absence (<100 nmol/mL) of the human milk oligosaccharide 2'-fucosyllactose. Information on secretor status was missing for two subjects.*

**AssetIndex2 = asset index; a proxy for socioeconomic status that was generated using the first principal component of a Principal Component Analysis (PCA) and reflects household wealth based on ownership of the following assets: electricity, radio, television, almirah, fan, table, chair, fridge, pump, freezer, phone, animals, mobile, watch, computer, autobike, bicycle, rickshaw and a vehicle[137]. Higher quintiles represent greater wealth. The lowest quintiles (Q1-Q2) and the highest quintiles (Q3-Q5) were grouped together for analysis.*

[§]*This binary variable was defined by self-reports collected at baseline. Households were categorized as 'treated' if they reported treating their drinking water (either using boiling and/or filtering) or 'untreated' if they reported no treatment of their drinking water. This variable was viewed as an indirect proxy for overall household hygiene and sanitation.*

[‖]*Reflects exclusively breastfed (only receiving human milk; no other liquids or solids, including water) versus not exclusively breastfed.*

[‡]*Reflects intra- and postpartum antibiotic usage by the mother.*

[#]*Reflects whether the infant had received any antibiotics up to the time of each stool sample collection timepoint.*
